# Cognitive and Neural State Dynamics of Story Comprehension

**DOI:** 10.1101/2020.07.10.194647

**Authors:** Hayoung Song, Bo-yong Park, Hyunjin Park, Won Mok Shim

**Affiliations:** Center for Neuroscience Imaging Research, IBS, Suwon, Korea; Department of Biomedical Engineering, Sungkyunkwan University, Suwon, Korea; Department of Psychology, University of Chicago, Chicago IL, USA; Department of Electronic, Electrical and Computer Engineering, Sungkyunkwan University, Suwon, Korea; McConnell Brain Imaging Centre, Montreal Neurological Institute and Hospital, McGill University, Montreal, QC, Canada; School of Electronic and Electrical Engineering, Sungkyunkwan University, Suwon, Korea

## Abstract

Understanding a story involves a constant interplay of the accumulation of narratives and its integration into a coherent structure. This study characterizes cognitive state dynamics during story comprehension and the corresponding network-level reconfiguration of the whole brain. We presented movie clips of temporally scrambled sequences, eliciting fluctuations in subjective feelings of understanding. An understanding occurred when processing events with high causal relations to previous events. Functional neuroimaging results showed that, during moments of understanding, the brain entered into a functionally integrated state with increased activation in the default mode network (DMN). Large-scale neural state transitions were synchronized across individuals who comprehended the same stories, with increasing occurrences of the DMN-dominant state. The time-resolved functional connectivities predicted changing cognitive states, and the predictive model was generalizable when tested on new stories. Taken together, these results suggest that the brain adaptively reconfigures its interactive states as we construct narratives to causally coherent structures.

## Introduction

We make sense of our own memory and others’ behavior by constantly constructing stories from an information stream that unfolds over time. Story understanding is a process of accumulating ongoing narratives, storing them in memory as situational contexts, and simultaneously integrating them to construct a coherent representation.^1,2^ Forming a coherent representation of a story involves understanding the causal structures of the events, including a logical flow of consecutive or temporally discontiguous events. Past research theorized that story understanding requires reinstating causally related past events and integrating them into a coherent narrative representation.^3,4^ However, empirical evidence is lacking on how the understanding of causal relations relates to the ongoing process of story comprehension.

Recent neuroscientific literature suggests that narratives are represented in activation patterns^5,6^ and functional connectivities (FCs)^7,8^ of the distributed regions in the default mode network (DMN), based on their capacity to integrate information over prolonged time periods.^9–11^ However, traditional cognitive models theorize that representation of narratives is updated by interaction of the broader networks of the whole brain, including regions involved in sensory processing, memory, and cognitive control.^12^ Yet, little is known about how large-scale functional networks dynamically reconfigure their representational and connective states as one’s story comprehension evolves over time. Prior research has indicated that large-scale brain networks alternate between functionally segregated and integrated states,^13–15^ depending on the information processing that is adaptively recruited at that moment.^16,17^ We hypothesize that a constant interplay of large-scale functional networks is crucial to story comprehension. Specifically, we assume that when narratives are actively integrated, individuals experience high levels of understanding, and an interactive state of functional networks of the brain emerges. In contrast, when individuals have low understanding and thus focus on accumulating available sensory and narrative inputs, we predict that the functional brain is biased toward a segregated state where each functional module operates independently.

Here, we characterized the cognitive processes involved in story comprehension and examined the dynamic reconfiguration of large-scale functional networks during story comprehension. To track cognitive state changes, we presented movie clips of temporally scrambled sequences and collected continuous behavioral responses when individuals experienced subjective feelings of understanding. Depending on the frequency of responses, we categorized time steps of the movies into moments of high or low levels of understanding. We separately measured causal relationships between pairwise moments of the movie and showed that moments of high understanding correspond to the moments when causally related past events are likely to be reinstated in memory. In functional neuroimaging experiments, we observed systematic modulations of blood oxygen-level dependent (BOLD) responses, as cognitive states alternated between the two distinct modes of narrative processing. On a large scale, the brain network enters into an integrated state during active comprehension, which is modulated by across-modular FCs of the default mode and frontoparietal networks. Using hidden Markov modeling (HMM) of latent neural states,^18^ we identified synchronized neural state dynamics across individuals when comprehending a novel story, with the DMN being a dominant state during high understanding moments. We further demonstrated that evolving cognitive states can be decoded from dynamic FCs of the whole brain, suggesting that FCs can be robust neuromarkers of distinctive modes of narrative information processing. By linking the cognitive and neural state dynamics of story comprehension, our study provides evidence that network-level reconfiguration underlies the ongoing naturalistic cognitive processes.

## Results

### Dynamic changes in subjective levels of story understanding

Three different silent films (10 min) were segmented into multiple scenes (36 ± 4 s per scene), and their temporal order was scrambled to induce fluctuations in participants’ subjective comprehension levels. In a behavioral experiment, subjects (N = 20 per film) watched the scrambled version of each film first (Initial Scrambled condition), and then the same film in a temporally intact sequence (Original condition). The scrambled order was constant for each film. To quantify the varying levels of understanding as they attempted to understand the stories, they were asked to press a button when they thought they had understood the story (“Aha”), or when their previous feeling of understanding turned out to be incorrect (“Oops”). As the “Oops” responses incorporate the psychological notion of “Aha”,^19^ no distinction was made between the two response types and were summed in the analysis. The results indicate that the moments of subjective understanding were largely consistent across individuals (Figure 1A). The responses of all subjects were summed with a sliding window of 36 s, in steps of 1 s. A window size of 36 s was chosen to match the average scene duration in the scrambled films. The aggregated responses were convolved with a canonical hemodynamic response function (HRF) to temporally relate to the functional magnetic resonance imaging (fMRI) responses. Based on the aggregated response frequency, the top one-third of the moments were categorized as moments of high understanding, and the bottom one-third were categorized as moments of low understanding (Figure 1B). We discarded the middle one-third of the moments because cognitive states during those moments were subject to higher variability across subjects than the top and bottom thirds.

**Figure 1.**
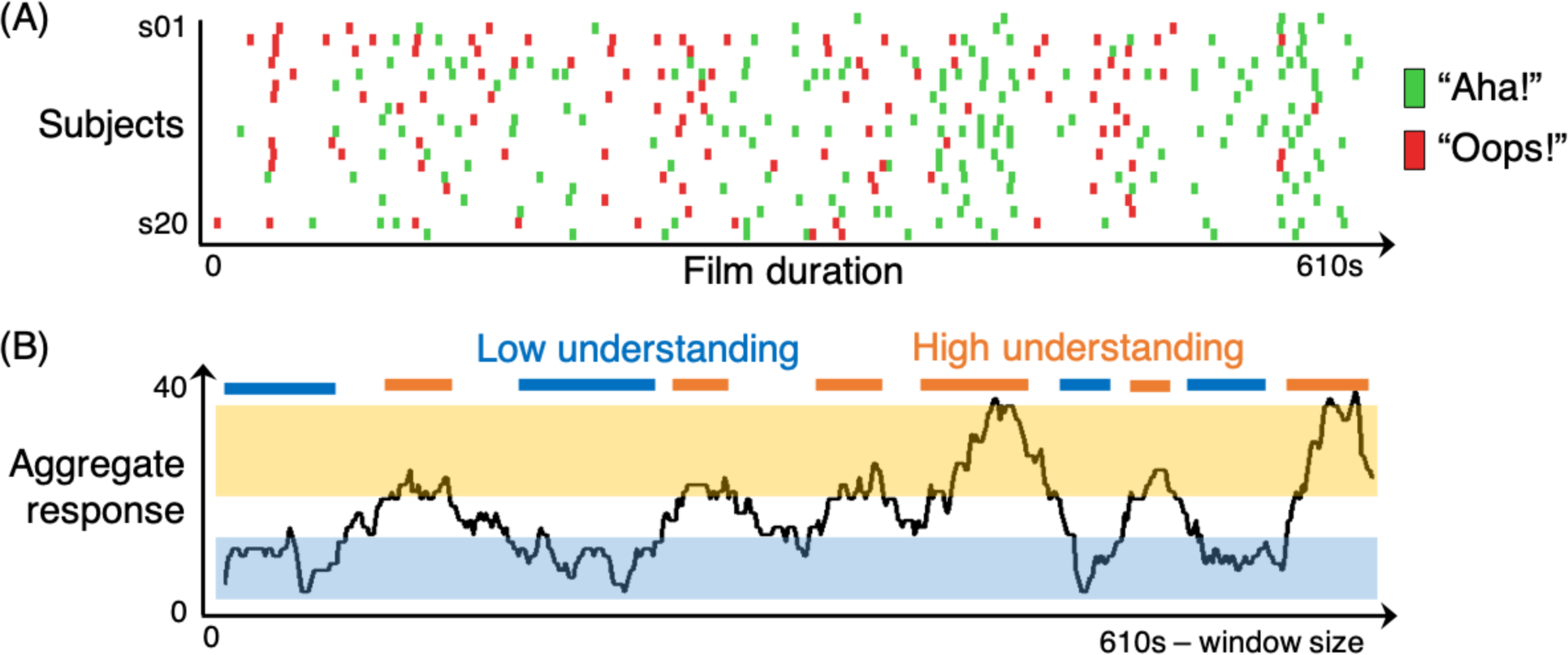
Dynamics of cognitive states during story comprehension. (A) Responses to moments of understanding while watching an exemplar scrambled film (N = 20). Moments of reported subjective understanding were consistent across subjects. The responses of all subjects were counted using a sliding time window. (B) Behavioral index of the changing level of understanding. The top one-third of the moments with frequent responses were defined as the moments of high understanding, whereas the bottom one-third were defined as the moments of low understanding.

### Reinstatement of the causally related past events during moments of understanding

We hypothesized that understanding narratives involves reinstating causally related past events to integrate with incoming information, thereby constructing a causally coherent representation of narratives.^3,20^ To test this hypothesis, we conducted a behavioral experiment (N = 12 per film) where subjects first segmented the events by marking the perceived event boundaries of the scrambled movie, then rated the causal relatedness of every possible pairwise event on a scale from 0 to 2: 0 (no causal relationship), 1 (shares a causal relationship), and 2 (shares a causal relationship that is critical in developing the story). The responses were summed to create moment-to-moment causal relationship matrices (Figure 2A), which displayed causal relationships that follow a logical flow of consecutive events as well as a causal chain between temporally discontiguous events (Figure 2B). We calculated causal relatedness of each moment by averaging the ratings of all preceding time points that were identified as being related to the current moment (Figure 2C). We predicted that the high understanding moments would correspond to moments with strong causal relations to past events. The level of understanding was highly correlated with the temporal changes in causal relatedness for all three films (Spearman’s *r* = 0.70, *r* = 0.40, *r* = 0.31, all *p*s < 0.001). Events corresponding to the high understanding moments had significantly higher causal relationships with past events than the events aligned to the low understanding moments (Wilcoxon signed-rank test; *z*(179) = 11.18, *z*(172) = 9.37, *z*(178) = 8.14; all *p*s < 0.001 for all films).

**Figure 2.**
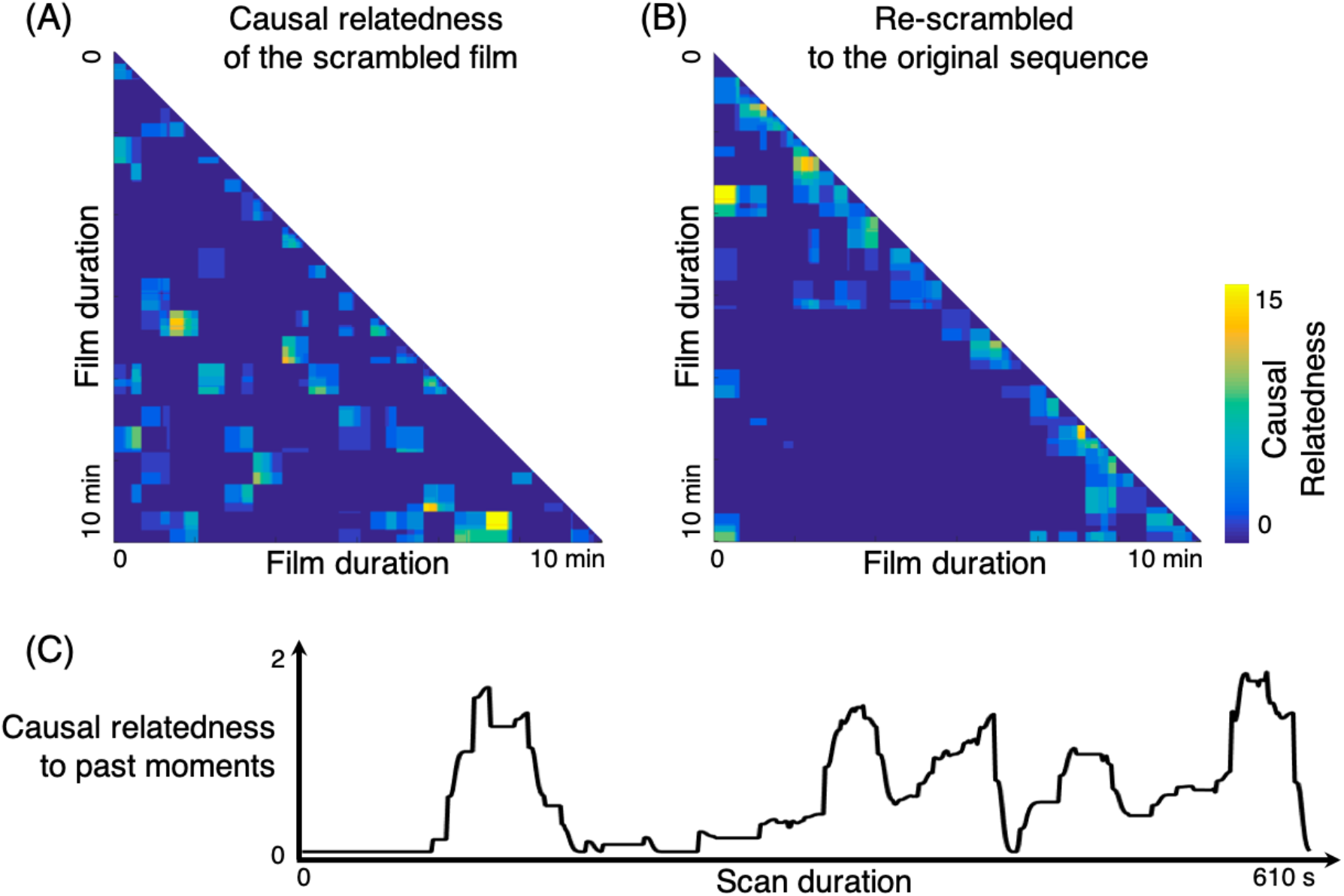
Causal relatedness between events in the films. (A) Causal relatedness matrix, indicating the causal relationships between all pairwise moments of an exemplar film (N = 12). Subjects rated the causal relatedness of the pairwise perceived event segments of the scrambled film on a scale from 0 to 2. All responses were summed to generate a single causal relatedness matrix. (B) Causal relatedness matrix, re-scrambled according to an original film sequence. Strong clustering around the diagonal indicates that the temporally contiguous moments in the original sequence tend to be causally linked. Pairwise events that are temporally distant but highly causally related also existed, indicating the presence of key events that are critical in developing narratives. (C) Degree of causal relatedness to past events. For each time point, we averaged the causal relatedness of every past moment (in different scenes) to the present moment. Moments of high causal relatedness to past events corresponded to the moments of high subjective levels of understanding, illustrated in Figure 1B.

There is a possibility that semantic relatedness between events, instead of causal relatedness, may have affected the comprehension of events. To test this alternative account, we measured semantic relatedness between pairwise moments of the films using the pairwise vector similarities derived from latent semantic analysis (LSA) of the sentences describing every moment of the event.^21^ We used the written annotations generated by native-level English speakers, giving detailed descriptions of each moment (2 s) in the films, including what was happening at that moment, by whom, where, when, how, and why. The words in every sentence were count-vectorized for LSA. Causal relatedness was significantly correlated with the semantic relatedness above the chance level (*p* < 0.001, *p* < 0.05, *p* < 0.001 for each film, compared to the null distribution of permuted semantic relatedness). Changes in understanding levels were positively correlated with changes in semantic relatedness (Spearman’s *r* = 0.32, *r* = 0.08, *r* = 0.25; all *p*s < 0.05), and the moments of high understanding had significantly higher semantic relatedness with the past compared to the moments of low understanding (*z*(179) = 8.10, *z*(172) = 3.52, *z*(178) = 6.33; all *p*s < 0.001). However, crucially, for all three films, the causal relatedness showed a significant relationship with the levels of understanding after the effect of the semantic relatedness was controlled for (partial correlations: *r* = 0.65, *r* = 0.44, *r* = 0.24; all *p*s < 0.001), whereas the semantic relatedness could not explain the understanding levels when the effect of causal relatedness was controlled for (*r* = 0.04, *r* = 0.08, *r* = 0.27; *p* = 0.29, *p* = 0.04, *p* < 0.001, respectively). The results suggest that reinstatement of the causally related past occurs during the high understanding moments, thereby constructing coherent representations of narratives.

### Modulation of BOLD responses

A separate, independent group of subjects underwent an fMRI experiment, where they watched the same sets of scrambled films inside a scanner (N = 24, 23, 20 for three film stimuli). As in the behavioral experiment, subjects underwent the Initial Scrambled and Original conditions, but additionally, they repeatedly watched the same scrambled film presented in the same order (Repeated Scrambled condition). To exclude possible task-induced effects, no explicit task was given, but subjects were instructed to attend to the stimulus and try to understand the original temporal and causal structure of the story. We examined if the BOLD responses were modulated by the changes in the level of understanding of the story. Using a general linear model (GLM), whole-brain BOLD responses were compared between the moments of high and low understanding based on the group-level behavioral index shown in Figure 1B.

In the Initial Scrambled condition, BOLD responses in the angular gyrus (Angular), precuneus (PreCu), posterior cingulate cortex (PCC), medial prefrontal cortex (mPFC), middle temporal gyrus (MTG), and middle frontal gyrus (MFG), which together comprise the DMN, showed higher levels of activity when a subject’s understanding was high. In contrast, when understanding was low, the frontal eye fields (FEF), a part of the dorsal attention network (DAN), and the visual sensory network including early and high-level visual areas showed higher levels of BOLD responses (Figure 3A). These results suggest that the DMN regions are involved when integrating narrative contexts to form an internal representation of the story, and that the DAN regions are involved when attending to a range of external sensory inputs that potentially convey narrative information. We predicted that the subjects remained attentive even during moments of low understanding, to accumulate necessary information for later comprehension. Similar modulations of BOLD responses were found when we applied the behavioral indices of understanding levels acquired while the original film was viewed during the Original condition of the fMRI dataset (Supplementary Figure S1), suggesting that similar modulations of the DMN and DAN occurred even when the film was viewed in a natural order.

**Figure 3.**
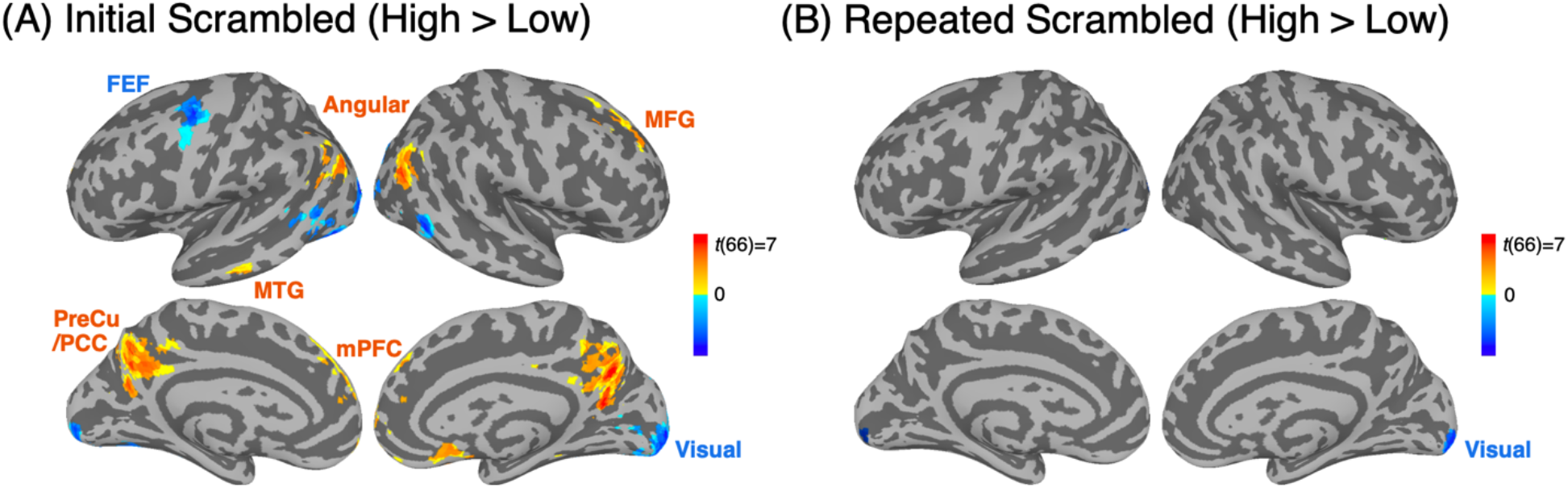
Contrasts in blood oxygen-level dependent (BOLD) responses between the moments of high and low understanding in the Initial and Repeated Scrambled conditions (cluster size > 40 and *q* < 0.01, N = 67). (A) Contrast between moments of high and low understanding, during the Initial Scrambled condition. When the understanding was high, responses in the default mode network increased, whereas responses in the dorsal attention network and visual sensory network decreased. (B) Contrast between the same moments during the Repeated Scrambled condition. No significant modulation of BOLD responses was observed, except decreased activation in a low-level visual region during high understanding. Detailed results of the general linear model analysis are summarized in Supplementary Table S1. Angular: angular gyrus, FEF: frontal eye fields, MFG: middle frontal gyrus, mPFC: medial prefrontal cortex, MTG: middle temporal gyrus, PreCu/PCC: precuneus/posterior cingulate cortex, Visual: visual cortex.

These results were also replicated when non-categorized, continuous behavioral indices of understanding levels were used as a regressor in a GLM (Supplementary Figure S2). Critically, in the Repeated Scrambled condition, none of the functional networks showed systematic modulation of BOLD responses, except for a small proportion of early visual areas, which showed greater activation during moments of low understanding, also found in the Initial Scrambled condition (Figure 3B; for a comparison between the Initial and Repeated Scrambled conditions, see Supplementary Figure S3). Since the identical stimulus was viewed in both the Initial and Repeated Scrambled conditions, a lack of difference between the high and low understanding moments during the Repeated Scrambled condition indicates that the results found in the Initial Scrambled condition were not driven by the intrinsic properties of the stimuli, but instead derived from the cognitive state differences that correspond to different levels of understanding. To examine stimulus-driven effects that may have resulted in larger BOLD responses in visual areas during low understanding moments, we assessed the physical salience of the movie frames by calculating the pixelwise stimulus intensities for every frame (1 s) of the movie. For all three films, salience was higher during moments of low understanding than during high understanding (Wilcoxon signed-rank tests, *z*(179) = 2.13, *z*(172) = 4.04, *z*(178) = 2.96 for each film; all *p*s < 0.05), suggesting that increased activity in the early-level visual areas is partially due to the higher stimulus intensities during moments of low understanding.

### Reconfiguration of dynamic functional connectivities and large-scale functional networks

We then examined if the whole brain reconfigures its FC patterns and large-scale functional networks as story comprehension evolves over time. During moments of low understanding, we expected a segregated state of the brain, where each functional network is engaged in its own specialized function. However, when integrating narratives into a coherent structure, we expected a tightly integrated state that enables efficient communication across distinctive functional systems^14,22^ For network analysis, we parcellated the brain into 122 regions-of-interest (ROIs)^23^ and grouped them into eight pre-defined functional networks, which includes the seven cortical functional networks of Yeo et al.^24^ – visual (VIS), somatosensory-motor (SM), DAN, ventral attention (VAN), limbic (LIMB), frontoparietal (FPN), and DMN – as well as one subcortical network (SUBC) consisting of subcortical regions extracted from the FreeSurfer segmentation (thalamus, striatum, hippocampus, and amygdala). To account for the dynamic changes in FCs in relation to cognitive state dynamics, we extracted the BOLD time course from each ROI and computed the time-resolved, regularized, and weighted FCs between all pairwise regions during the Initial Scrambled, Original, and Repeated Scrambled conditions, respectively (Figure 4A). Graph theoretical measures were computed in a time-resolved manner to capture changes in large-scale functional network structures.^25^ To examine the degree of functional segregation, we measured modularity, which is the degree to which functionally specialized regions of the brain are clustered in a modular structure.^26^ As an indicator for functional integration, we measured global efficiency that reflects integrative information processing across remote regions of the brain.^16,27,28^

**Figure 4.**
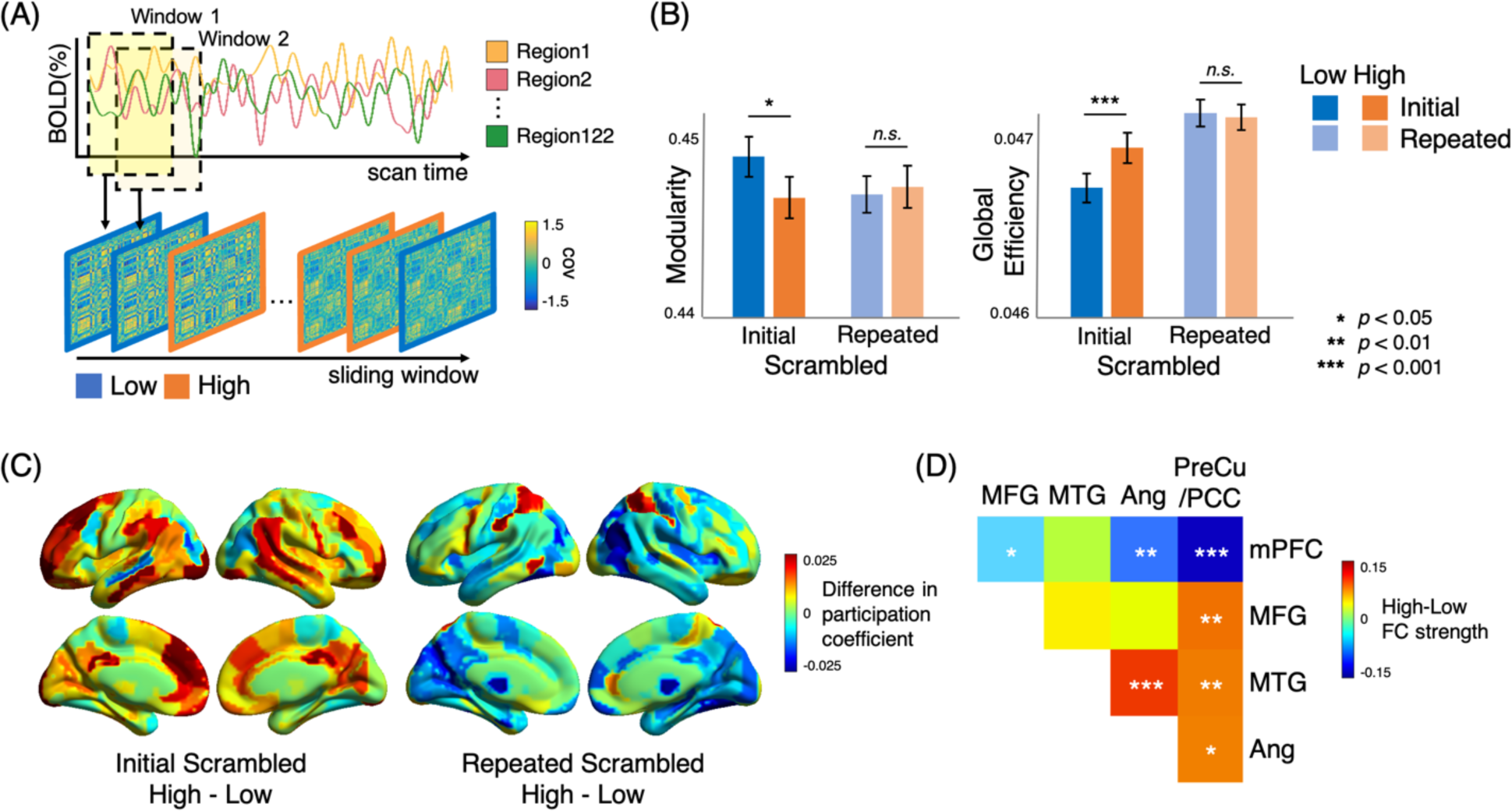
Dynamic reconfiguration of large-scale functional networks. (A) Schematic overview of dynamic functional connectivity (FC) analysis using a sliding window.^29^ The blood oxygen-level dependent (BOLD) time series was extracted from 122 regions of interest (ROIs).^23,24^ Time-resolved FC matrices were constructed for each window across film duration, and graph theoretical network measures were computed. Network measures for each moment were categorized by their correspondence to the cognitive states of either high or low understanding. (B) Global network reconfiguration corresponding to cognitive state changes. During moments of high understanding, the brain showed a functionally integrated state with lower modularity and higher efficiency. The difference was observed in the Initial Scrambled but not Repeated Scrambled condition. (C) Regional network reconfiguration dependent on the level of understanding. The difference in participation coefficients of 246 ROIs between the moments of high and low understanding, averaged across subjects. We used the ROIs defined in the Brainnetome atlas^30^ to ensure whole-brain coverage, and visualized them with BrainNet Viewer (https://www.nitrc.org/projects/bnv/).^31^ (D) Within-network FC strengths of the pairwise subregions of the DMN, dependent on the level of understanding. The colors indicate network strength differences between moments of high and low understanding, averaged across subjects. When the understanding was high, FCs of the mPFC with the rest of the regions in the DMN decreased, whereas the FCs of the PreCu/PCC with the rest of the DMN regions increased except with mPFC. mPFC: medial prefrontal cortex, PreCu/PCC: precuneus/posterior cingulate cortex.

In the Initial Scrambled condition, modularity decreased when the understanding was high (Wilcoxon signed-rank test on the combined data of three different films; *z*(66) = 2.32, *p* < 0.05), suggesting that tight interconnections across different functional modules arise when information is being integrated into coherent narratives (Figure 4B). In contrast, there was no difference in modularity between high and low understanding moments during the Repeated Scrambled condition (*z*(66) = 0.58, *p* = 0.561). A significant interaction was found between the viewing conditions (Initial and Repeated) and the understanding levels (high and low; *F*(1,66) = 6.98, *p* < 0.05), although no main effect was observed (all *p*s > 0.1). Similarly, global efficiency was higher during moments of high understanding compared to low understanding (*z*(66) = 3.67, *p* < .001). There was no difference in global efficiency during the Repeated Scrambled condition (*z*(66) = 0.58, *p* = 0.566), and the interaction was significant (*F*(1,66) = 10.62, *p* < 0.01). Notably, a significant main effect of the Scrambled conditions was found (*F*(1,66) = 26.27, *p* < 0.001), with a higher efficiency when the same scrambled film was watched repeatedly. These results suggest that the efficiency of information processing increases when a coherent situational context is represented in the brain, consistent with previous findings of enhanced FCs within the DMN when scrambled movies were watched repeatedly.^7,8^ The results were robust when different sizes of sliding window or a different cortical parcellation were used (Supplementary Figure S4).

Along with a global reconfiguration, the time-resolved regional network measures between the moments of high and low understanding were compared (Figure 4C). For all ROIs, the participation coefficient and within-modular degree z-score were measured, which indicate the degrees of across-modular and within-modular connections, respectively. A higher participation coefficient indicates that a region is functionally connected to the regions of other functional networks in a distributed manner, whereas a higher within-modular degree z-score indicates that a region is mainly associated with the regions that lie within the same network.^32^ During the Initial Scrambled condition, all functional networks showed higher participation coefficients when the understanding was high compared to low. In particular, the FPN (*z*(66) = 4.24, FDR-*p* < 0.001, corrected for multiple comparisons of functional networks) and DMN (*z*(66) = 2.87, FDR-*p* < 0.05) showed significantly higher participation coefficients for high understanding compared to low understanding (Figure 4C). There was a significant interaction between the Scrambled conditions and understanding levels for both networks (FPN: *F*(1,66) = 11.93, DMN: *F*(1,66) = 12.29; both *p*s < 0.001; Supplementary Figure S5A). During the Repeated Scrambled condition, no functional network showed similar patterns of modulation in their across-modular FCs, except for a reversed pattern of higher participation coefficients during low understanding in VIS and SUBC (*z*(66) = 3.35, *z*(66) = 3.21 respectively; both FDR-*p*s < .01). The within-modular FCs, quantified by within-modular degree z-scores, did not differ across the moments of high and low understanding for any of the functional networks, during both the Initial and Repeated Scrambled conditions (all FDR-*p*s > 0.6; Supplementary Figure S5B). The results were replicated with different cortical parcellation (Supplementary Figure S6). Overall, these results indicate that the brain enters into a functionally integrated state that ensures more efficient information transfer across functional modules when integrating narratives to coherent representation. Such functional integration is driven by the increased across-modular FCs of the FPN and DMN, but not by the within-modular FCs.

Although the distributed regions of the DMN are tightly associated through functional connections and share common patterns of deactivation during tasks, previous literature has discussed how the regions within the DMN take on different functional roles.^33,34^ To specify the regional roles within the DMN during story comprehension, we compared the FC strength of the pairwise subregions of the DMN between the moments of high and low understanding (Figure 4D). Notably, the mPFC showed decreased FCs with all other regions in the DMN (all FDR-*p*s < 0.05, except for a non-significant but decreasing trend of FC with MTG), whereas the PreCu/PCC showed increased FCs with all other DMN subregions except its connection to the mPFC (all FDR-*p*s < 0.05) when the understanding was high. Increased FCs of the PreCu/PCC and decreased FCs of the mPFC, paired with other subregions of the DMN during the moments of high understanding, support distinct functional roles of the anterior and posterior subregions of the DMN.^33.34^

### Latent neural state dynamics corresponding to changes in cognition

We investigated whether low-dimensional neural state dynamics track changes in cognitive states involved with story comprehension. To infer the dynamics of latent states in an unsupervised data-driven manner, we applied the HMM, which assumes that the observed sequence of brain activity is probabilistically conditioned on the sequence of discrete latent states.^35–38^ To characterize the observed sequence of neural activities, we first conducted group-level independent component analysis (ICA)^39^ from all subjects’ concatenated fMRI responses of all three conditions across three different films. Thirty independent components (ICs) were identified to be signal components that were involved during film watching. The discrete latent neural states were derived from patterns of activation and functional covariance of the 30 ICs, as we trained the HMM on the data from the Original condition. When we set the number of latent states to four (for information regarding the choice of the optimal number of states, see Supplementary Figure S7), the extracted states were: i) SM + VIS, ii) DAN, iii) Integrated DMN + VIS, and iv) Segregated DMN + VIS (Figure 5A). Each state was labeled as one or the combination of eight functional networks that showed the highest levels of BOLD activation (Supplementary Table S2). Notably, the two DMN + VIS states, (iii) and (iv), showed indistinguishable activation patterns, yet their functional covariance significantly differed such that one had higher across-modular FCs and lower within-modular FCs than the other (all FDR-*p*s < 0.05). The one with higher across-modular and lower within-modular FCs was termed the “Integrated” DMN + VIS state, and the other the “Segregated” DMN + VIS state. We applied the derived states to infer the latent state dynamics in the Initial and Repeated Scrambled conditions. First, the inferred latent states were verified to be dynamic in nature. The maximal fractional occupancy, the highest proportion of a particular state’s occurrence across all time points per subject, was below 50% for most of the subjects (*p*s < 0.001; Supplementary Figure S8), indicating transitions from one latent state to more than one other state.^35^

**Figure 5.**
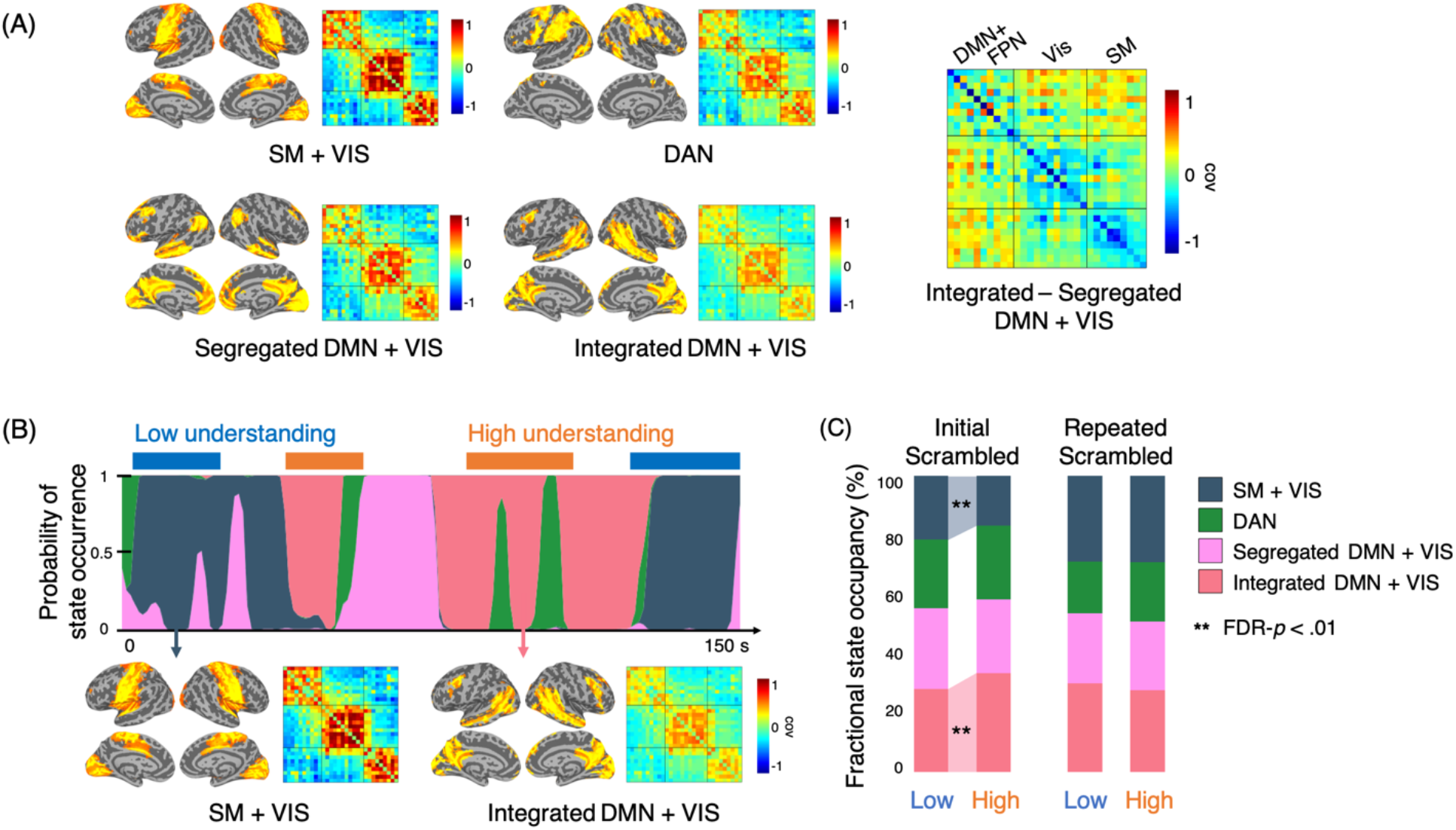
Latent neural state dynamics inferred using the hidden Markov model (HMM). (A) Activation patterns and functional covariance of the four latent states, identified by training the HMM with the Original condition. The SM + VIS, DAN, Segregated DMN + VIS and Integrated DMN + VIS were termed based on their levels of blood oxygen-level dependent (BOLD) activation corresponding to the eight pre-defined functional networks. The covariance matrix on the right shows the difference between Integrated DMN + VIS and Segregated DMN + VIS. Integrated DMN + VIS is characterized by higher across-modular connections and lower within-modular connections compared to Segregated DMN + VIS. (B) State occupancy and transition dynamics of a representative subject during the Initial Scrambled condition. Occurrences of the states were probabilistically inferred for each time point.^35,36^ (C) The average fractional occupancy of the four latent states during the moments of high and low understanding, in the Initial and Repeated Scrambled conditions. The highlighted background between the colored bars indicates significant differences in fractional occupancy. In the Initial Scrambled condition, Integrated DMN + VIS occurred more frequently during high understanding, whereas SM + VIS emerged more frequently during low understanding. No modulations in state occupancy were observed in the Repeated Scrambled condition. DAN: dorsal attention network, DMN: default mode network, FPN: frontoparietal network, SM: somatosensory-motor networks, VIS: visual network.

Next, we examined whether the fractional occupancy of each neural state was modulated as the cognitive states traversed between different levels of understanding. Figure 5B illustrates the dynamics of the state occurrence probabilities of an exemplar subject, which is mapped in time with the behavioral index of understanding levels. In the Initial Scrambled condition, the SM + VIS had a higher occupancy during the moments of low understanding (*t*(66) = 2.84, FDR-*p* < .05), whereas the Integrated DMN + VIS had a higher occupancy during the moments of high understanding (*t*(66) = 3.64, FDR-*p* < .01) (Figure 5C). None of the states differed in fractional occupancy across the two understanding levels in the Repeated Scrambled condition (all FDR-*p*s > 0.2). Consistent results were found when six latent states instead of four were used in the HMM (Supplementary Figure S9A-C). These results imply that the DMN, in tight connection with sensory networks, is highly involved when the narratives are actively integrated. In contrast, when one focuses on accumulating information from external inputs, the low-level sensory and motor networks take over its role (for a comparison of fractional occupancy between the Scrambled conditions, see Supplementary Figure S10).

Furthermore, we investigated whether the neural state dynamics were synchronous across individuals as they comprehended the same stories. Figure 6A illustrates the inferred neural state sequence for all subjects as they watched an exemplar film stimulus in the Initial and Repeated Scrambled conditions. The proportion of the moments when the inferred state was identical was calculated for all pairwise subjects per film. The neural state dynamics were more synchronized across subjects in the Initial than in the Repeated Scrambled condition (paired t-test, *t*(718) = 22.86, *p* < 0.001, Cohen’s *d* = 1.04; Figure 6B), which was replicated when six states were used (Supplementary Figure S9D). These results suggest that individuals share similar neural dynamics when actively trying to understand a novel story, yet the synchrony decreases when the story is no longer novel due to idiosyncratic transitions of the cognitive or attentional states in each individual.

**Figure 6.**
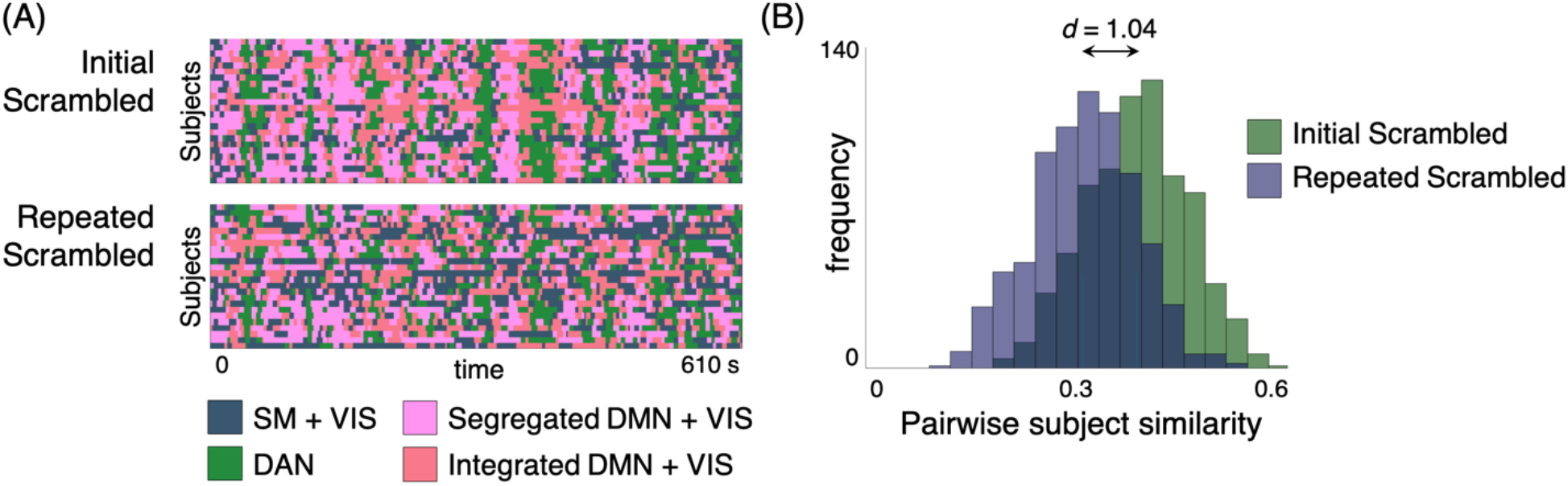
Synchrony of the latent neural states across individuals during the Initial and Repeated Scrambled conditions. (A) The neural state dynamics of all subjects in the Initial (top) and Repeated Scrambled (bottom) conditions for an exemplar film. (B) Histogram of the similarities in state dynamics for all pairwise subjects in the two conditions. The neural state dynamics were more similar across individuals in the Initial compared to the Repeated Scrambled condition, indicating that common neural states emerge when individuals are actively trying to understand a story, compared to moments when they repeatedly watch the same story. DAN: dorsal attention network, DMN: default mode network, FPN: frontoparietal network, SM: somatosensory-motor networks, VIS: visual network.

### Prediction of the evolving cognitive states using time-resolved functional connectivity signatures

We examined whether a subjective level of understanding can be predicted from the neural signatures. Compared to traditional predictive modeling, where a single pattern of brain activity is linked to a trait-level behavioral score for each subject,^40^ we conducted dynamic predictive modeling, which maps time-resolved brain patterns to the time-resolved behavioral indices (Figure 7A). The time-resolved brain patterns were measured by applying a sliding window analysis on the FC matrices or the regional BOLD time courses, and the dynamic behaviors were represented by the binary index of high versus low understanding at each moment, collected from the independent behavioral study (Figure 1B). We trained a linear support vector machine (SVM) to decode moments of high or low understanding, given the multivariate neural features at each time point. The model was cross-validated in a leave-one-subject-out fashion, either on data within the same film (within-film decoding) or across different films (across-film decoding; Figure 7B). The within-film decoding procedure was cross-validated across subjects who watched the same film. The across-film decoding was conducted by training the model on the data collected from two of the three films and testing on a held-out subject who watched the held-out film. We performed the across-film decoding to exclude the possibility that stimulus-driven properties, other than the cognitive states associated with story understanding, inflate the decoding performance. Moreover, we compared the performance of the predictive models using two types of neural signatures - the time-resolved FC strength of the pairwise ROIs and the activation patterns of each ROI - both spanning the whole brain. An additional feature selection procedure was employed in the FC pattern-based decoding. The pairwise regions of which time-resolved FCs were consistently correlated with the continuous behavioral index of story understanding were selected in every cross-validation fold (*p* < 0.01; for the number of selected features, see Supplementary Table S3).^40^ Feature selection was not applied in the activation pattern-based decoding due to its initial small number of features. The pairwise FCs consistently selected (>80% of cross-validation folds) in the across-film decoding are shown in Figure 7C. The number of FCs consistently selected across folds was far larger in the Initial (208 pairs, 2.8% of the total pairs) than in the Repeated (3 pairs) Scrambled condition, suggesting that the brain regions modulate their neural responses synchronously with other regions as one actively constructs coherent narratives. In particular, the FPN and the two attentional networks, DAN and VAN, were selected above the chance level compared to a null distribution, where the network indices were permuted while preserving the total number of features (all FDR-*p*s < 0.001).

**Figure 7.**
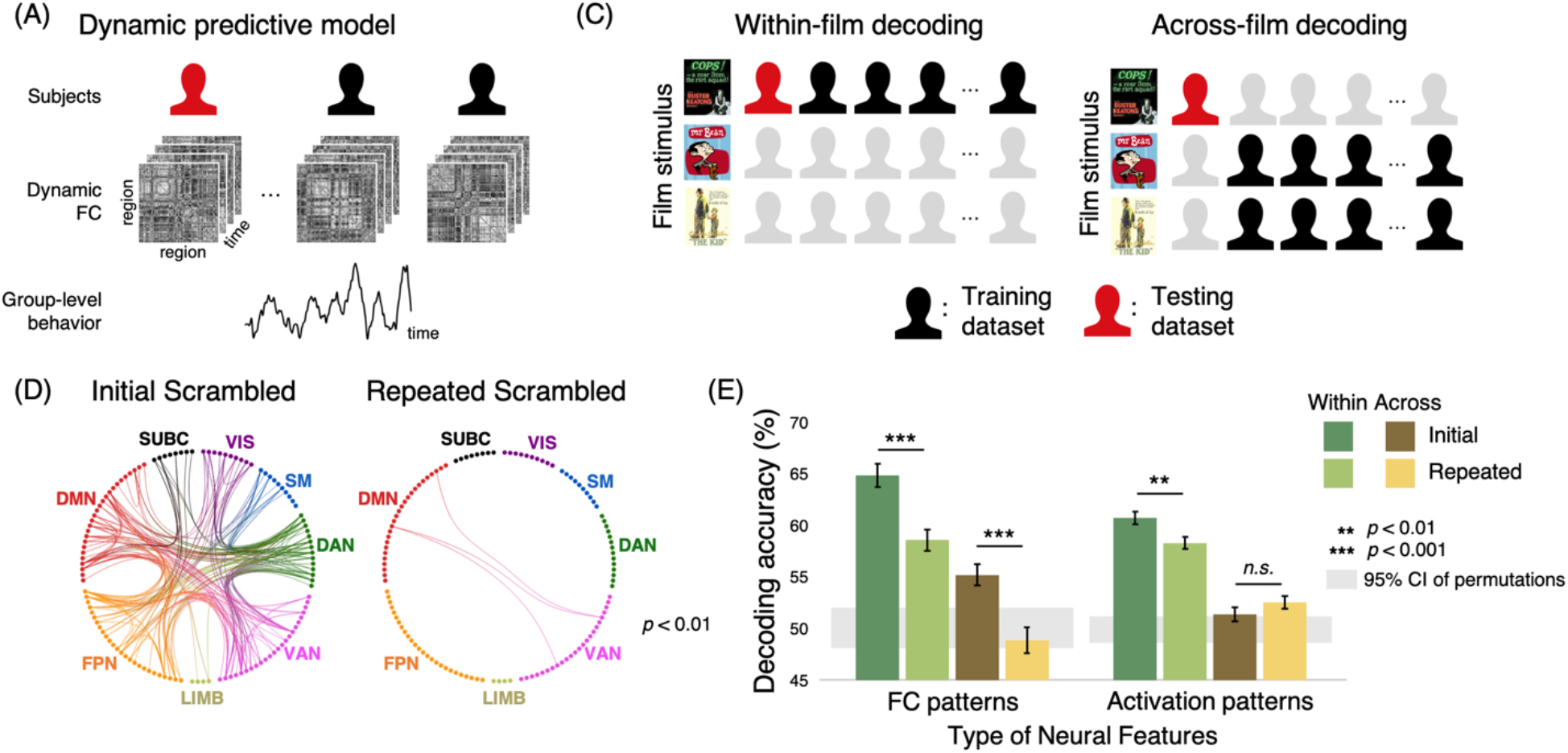
Prediction of the moment-to-moment cognitive states, using the functional connectivities (FCs) and activation patterns of the whole brain. (A) Schematics of dynamic predictive modeling. A traditional predictive model learns mapping between individual’s brain patterns and corresponding behavioral scores in training datasets, to predict held-out individual’s behavior.^40^ An individual subject is represented by a single brain pattern and a trait-level behavioral score, which discounts temporal variability. A dynamic predictive model, however, predicts time-resolved behaviors (e.g., changes in behavioral performance or changes in cognitive states) from time-resolved brain patterns. (B) Illustration of the two cross-validation schemes. In the within-film decoding, the model was tested on the held-out subject who watched the same film. In the across-film decoding, the model was tested on the held-out subject who watched the held-out film. (C) The pairwise regions that were consistently selected as features (>80% of cross-validation folds) in the across-film decoding. A larger number of FCs were selected to be significantly correlated with the changes in cognitive states during the Initial compared to the Repeated Scrambled condition. (D) Prediction performance. Although both FC- and activation-pattern based predictions were successful in decoding the cognitive state in the within-film decoding, only the FC pattern-based prediction was successful in the across-film decoding. The gray areas indicate 95% confidence intervals of the decoding performance using permuted behavioral indices. DAN: dorsal attention, DMN: default mode network, FPN: frontoparietal network, LIMB: limbic network, SM: somatosensory-motor networks, SUBC: subcortical networks, VAN: ventral attention network, VIS: visual network.

The results of the predictive modeling are illustrated in Figure 7D. In the within-film decoding, the predictive performance was higher for the Initial than for the Repeated Scrambled condition, when either FC patterns (*z*(66) = 4.30, *p* < 0.001) or activation patterns (*z*(66) = 2.95, *p* < 0.01) were used. A significant interaction between the Scrambled conditions (Initial and Repeated) and the type of neural features (FC patterns and activation patterns) was found (*F*(1,66) = 7.78, *p* < 0.01), which indicates a larger performance improvement in the Initial compared to the Repeated Scrambled condition when FC patterns were used for decoding. In the across-film decoding, when the FC patterns were used, the prediction accuracy of the Initial Scrambled condition outperformed that of the Repeated Scrambled condition (*z*(66) = 3.43, *p* < 0.001). However, when activation patterns were used, the performance did not differ between the Initial and Repeated Scrambled conditions (*z*(66) = 1.40, *p* = 0.161), leading to a significant interaction (*F*(1,66) = 17.43, *p* < 0.001). The superior performance of the FC pattern-based across-film decoding suggests that the FCs of the whole brain contain information on cognitive states that can generalize across stories. All conditions, except FC pattern-based, across-film decoding in the Repeated Scrambled condition (*p* = 0.828) were significantly different from a chance-level distribution (1,000 permutations of behavioral indices, all *p*s < 0.05).

The results may have been driven by the feature selection procedure, which was only applied to the FC pattern-based decoding. Thus, we used the activation patterns of the whole brain voxels instead of the ROIs and applied the same feature selection method as the FC pattern-based decoding (Supplementary Figure S11). The results remained consistent, which indicates that the failure of the across-film decoding using activation patterns (Figure 7D) cannot be attributed to either the smaller number of features or the absence of feature selection. To examine if specific functional networks selectively contribute to decoding cognitive states, we conducted decoding analyses using the FCs of each functional network separately as a seed (Supplementary Figure S12A), or the activation patterns of the ROIs within respective functional networks (Supplementary Figure S12B). The trend of results obtained from the whole brain was comparable when respective functional networks were tested individually, indicating that the cognitive states related to one’s level of story understanding are not restricted to the operation of a particular functional system. Overall, the results revealed that the cognitive states associated with story understanding are more robustly represented by the dynamic FCs that span the whole brain, as opposed to regional activation patterns.

## Discussion

This study characterizes story understanding as the dynamic interplay of information accumulation (i.e., when having low understanding of narratives) and integration (i.e., when active understanding occurs). We identified the dynamic fluctuation of these cognitive states rom subjects’ behavioral responses as they watched temporally scrambled films (Figure 1). By assessing the causal relationships of the events, we showed that the moments of narrative integration corresponded to the moments when causally related past events were likely to be reinstated in memory (Figure 2). Using fMRI, we showed how large-scale functional networks of the brain adaptively reconfigure its activation and connective states when individuals engage in comprehending narratives. The systematic modulation of BOLD responses was observed, with higher DMN activities during moments of high understanding, and higher DAN activities during moments of low understanding (Figure 3). Additionally, whole brain network-level reconfiguration was aligned to cognitive state changes. The networks entered a functionally integrated state during moments of high understanding, supported by the across-modular connections of the DMN and FPN (Figure 4). Using HMM, we showed that the DMN in a functionally integrated state becomes dominant during information integration, whereas sensory and motor networks become dominant during information accumulation (Figure 5). The latent states were synchronized across subjects when comprehending novel stories (Figure 6). We further demonstrated that evolving cognitive states of unseen individuals can be predicted using time-resolved FCs, and that the prediction can be generalized across different stories (Figure 7).

Prior fMRI studies on large-scale networks focused on comparing brain states during task versus rest,^15^ tasks with different cognitive loads,^14,41^ or focused versus unfocused attentional states.^42,43^ However, dynamic state changes that accompany cognition in naturalistic settings are far more complex and mostly occur without explicit task structures. Understanding a temporally scrambled story necessitates a constant attentional focus, but at the same time requires continuous shifts in cognitive states between information accumulation and integration. Previous work theorized the notion of ‘external’ and ‘internal’ modes of information processing, illustrating how the brain undergoes state changes between accumulating the information from the external world, and integrating the accumulated information with a focus on internal, self-generated thoughts.^44,45^ We suggest that story comprehension is an exemplary naturalistic cognitive process that entails adaptive switching between the two states of information processing. Our study further illustrates that dynamic switches between segregated and integrated states of functional networks of the brain, which were previously suggested to reflect the efficiency and flexibility when performing cognitive tasks or during rest,^14,16,17,41^ are also associated with the externally- and internally-directed modes of information processing during story comprehension.

Our results are consistent with previous findings indicating DMN as a hub of narrative information processing, with its activation patterns representing discrete events,^5,6^ and its within-network FCs representing a degree of story coherence.^7,8^ Although encoding of events may be mediated by the regional BOLD activation patterns, our results imply that the process of constructing narratives by integrating inputs with one’s memory is likely to be mediated by the synchronous or de-synchronous activities across distinct functional networks. Story comprehension may require collaborative and collective operations of large-scale networks, which efficiently reconfigure their network states depending on the type of cognitive processes adaptively recruited at that moment.

Future studies are necessary to investigate the underlying computational mechanisms explaining the accumulation and integration of narratives^46,47^ in relation to the neural representation formed on a moment-to-moment basis.^48–50^ Our work provides insights into the cognitive and computational processes on how representations of discrete events across distant times in memory, chained with causal relations, may be integrated with accumulated inputs in real-time to form coherent narratives. Communication of information between functional modules of the brain during story comprehension may be implemented via modulation of FCs and global reconfiguration of the dynamic brain states.

## Acknowledgements

We thank Youngmin Jeon for assistance with behavioral data collection, and Boohee Choi for technical support in fMRI data collection. We thank Jeongjun Park, Hun Seok Choi, Woochul Choi, and Monica D. Rosenberg for constructive feedback on the manuscript and thank Oliver James, Choong-wan Woo, and Janice Chen for their feedback on data analyses. This research was supported by the Institute for Basic Science Grant (IBS R015-D1) and the National Research Foundation (NRF) of Korea Grant by the Korean government (NRF-2019M3E5D2A01060299 and NRF-2019R1A2C1085566).

## Author Contributions

Conceptualization, H.S., and W.M.S; Methodology, H.S., B.-Y.P., H.P., and W.M.S.; Formal Analysis, H.S.; Investigation, H.S., and W.M.S.; Writing - Original Draft, H.S.; Writing - Review & Editing, H.S., B.-Y.P., H.P., and W.M.S.; Funding Acquisition, W.M.S.; Supervision, W.M.S.

## Declaration of Interests

The authors declare no competing interests.

## Methods

### Subjects

Independent groups of subjects participated in two behavioral and one fMRI experiments (Behavioral Experiment 1: 20 subjects per film, with a total of 27 subjects; five women, mean age 22.6 ± 2.1 years. Behavioral Experiment 2: 12 subjects per film, with a total of 29 subjects; 10 women, mean age 21.6 ± 2.2 years. FMRI Experiment: 24 subjects for *Cops*, 23 subjects for *The Kid*, and 20 subjects for *Mr. Bean*, with a total of 30 subjects; 10 women, mean age 24 ± 2.1 years). None of the subjects had watched the films prior to the experiment. All but one subject were native Korean speakers. All subjects who participated in the fMRI study were right-handed, except one. Subjects reported no history of visual, hearing, or any form of neurological impairment. The subjects provided informed consent before taking part in the study and were monetarily compensated. The study was approved by the Institutional Review Board of Sungkyunkwan University.

### Film Stimuli

Three movie clips - *Cops* (1922, Keaton & Cline), *The Kid* (1921, Chaplin), and *Mr. Bean: The Animated Series, Art Thief* (season 2, episode 13; 2003) - were used in both behavioral and fMRI experiments. Since the films did not contain any form of verbal communication or narration, narratives were mainly delivered in the visual modality. Each original film was edited to a 10 min version where a coherent narrative was complete within the given time. Each film was divided into 16 (*The Kid*) or 17 (*Cops* and *Mr. Bean*) scene segments and shuffled in pseudorandom order, such that dynamic changes in the subjective level of understanding could be induced. Most segments were demarcated by the original director’s cut, and the length of each segment was adjusted to between 32 s and 40 s, with 1 s increments. The speed adjustment was minor, and none of the subjects reported perceived differences in the speed of the scenes.

### Behavioral Experiment 1: Subjective level of understanding

We collected behavioral responses while subjects were watching a film in a scrambled sequence (Initial Scrambled) and then in the original sequence (Original). The stimuli were presented by GStreamer library (Open-Source, 2014) and the responses were recorded with Matlab (Mathworks, Natick, MA, USA) and Psychtoolbox.^51,52^ The experiment was conducted in a dimly lit room where the films were presented on a CRT monitor. Prior to the experiment, subjects participated in a practice session with a different movie clip, *Oggy and the Cockroaches: The Animated Series, Panic Room* (season 4, episode 8; 2013). Subjects pressed a button when they thought that they had understood the story (“Aha” response: e.g., when the temporal sequence or the causal relationship of the original story were understood, or when interim understanding of previously presented events occurred), and pressed another button when they realized their prior understanding was incorrect (“Oops” response). The experiment started with the Initial Scrambled condition, followed by a 30 s rest with a blank screen, then followed by the Original condition. After watching the films, subjects completed a comprehension quiz about the plots and contents of the story. The quiz consisted of six true or false questions, four short answer questions, and two questions on the original temporal sequence of the film. The data of two subjects who scored exceptionally low were excluded from the analyses. The button responses of both “Aha” and “Oops” of all subjects were summed without distinction in a sliding window of 36 s with a step size of 1 s. The two types of responses were not discerned in the analysis, because the psychological notion of “Oops” contains the state of “Aha.”^19^ We initially chose a window size of 36 s to match the average duration of each scene segments in the scrambled films, but replicated our results with window sizes of 24 s, 30 s, 36 s, and 42 s. The number of aggregated responses at each moment was convolved with canonical HRF to relate to the BOLD responses. The top one-third of the moments with frequent responses across subjects were labeled as the moments of ‘high understanding’ and the bottom one-third were labeled as the moments of ‘low understanding.’ We discarded the middle one-third due to low consistency in responses across subjects. With an assumption that cognitive states of understanding are not instantaneous but prolonged, the labeled moments were discarded if they did not persist for at least 10 consecutive time points. The number of discarded time points was small; 6.02 ± 5.72% of the one-third splits of the total time points.

### Behavioral Experiment 2: Degree of causal relatedness to past events

We collected reports of causal relations between all pairwise events of the scrambled films. The stimuli presentation and response recording were controlled with Adobe Premiere Pro CC (Adobe Systems, San Jose, CA, US). Subjects initially watched the film in scrambled and original orders. Then, they were asked to segment the scrambled film in terms of the events’ narrative contents, by marking event boundaries without limit on the total number as they freely swiped through the film on the video editor.^6,53^ The scene boundaries created by initial temporal scrambling were marked in the video editor. Next, subjects were instructed to write a short description of each segmented event. Finally, using the descriptions of the events, subjects rated the degree of causal relatedness of all possible pairwise events on a scale of 0 to 2. A pair of events was rated 1 if one event was causally attributed to, or explained by, the happening of another event. A pair was rated 2 if a causal relatedness between pairwise events played a main role in developing the key narratives of the story. The pairwise events that had no significant causal relatedness were rated 0. Subjects performed the task at their own pace without time limit. The timings of the perceived event boundaries were rounded to a 1 s sampling rate. By summing all the subjects’ ratings, a causal relatedness matrix for each film was constructed, which specified degrees of causal relatedness of all pairwise moments for the duration of the film. We re-scrambled the event sequence back to the original order to validate the key causal relatedness of events that were critical for story development. The causal relatedness score of each moment was computed by averaging every past moments’ causal relatedness to the present moment. High causal relatedness scores indicated that the past events were more likely to be reinstated in memory while a current event was processed.

### Control Experiment: Degree of semantic relatedness with past events

As a control experiment, we measured the degree of semantic relatedness between all pairwise events of the scrambled films. Written annotations of the narrative contents were given to every 2 s of the scrambled films by four annotators (four females, mean age 24.5 ± 1.3 years) with native-level English proficiency, including the first author. The annotators had never watched the films prior to the annotations, except the first author. The example annotations in Nishida & Nishimoto (2018) were used to instruct the annotators.^54^ Specifically, the annotators were instructed to make detailed descriptions of each scene, including what is happening at the current moment, by whom, where, when, how, and why. All words included in every 2 s of the annotations were count-vectorized. Latent semantic analysis (LSA; sklearn.decomposition.TruncatedSVD)^21^ was conducted, which quantifies the semantics of every 2 s annotation by the distribution of words using singular value decomposition (dimensionality set to 100). The semantic relatedness between the pairwise moments of the films were calculated by the cosine similarities between the LSA output vectors. The semantic relatedness score of each moment was computed by averaging every past moments’ semantic relatedness to the present moment.

### Control Experiment: Stimulus saliency

The visual salience was measured for all video frames at a sampling rate of 1 s. The pixelwise intensity of each frame was measured by SaliencyToolbox,^55^ and the intensities of every location was averaged to represent framewise salience. The saliency measures of the frames that corresponded to the moments of high and low understanding were compared using a Wilcoxon signed-rank test.

### Functional MRI experiment

Subjects were scanned with a 3T scanner (Magnetom Prisma; Siemens Healthineers, Erlangen, Germany) with a 64-channel head coil. A session consisted of one anatomical run and one task-based functional run. The anatomical images were acquired before or after viewing the film using T1-weighted magnetization-prepared rapid gradient echo pulse sequence (repetition time (TR) = 2,200 ms, echo time (TE) = 2.44 ms, field of view = 256 mm × 256 mm, and 1 mm isotropic voxels). Functional images were acquired using a T2*-weighted echo planar imaging (EPI) sequence (TR = 1,000 ms, TE = 30 ms, multiband factor = 3, field of view = 240 mm × 240 mm, and 3 mm isotropic voxels, with 48 slices covering the whole brain). A single EPI run lasted for 31 m 20 s, which included 30 s of blank fixation periods in between the three film-watching conditions (Initial Scrambled, Original, and Repeated Scrambled), and 10 s of additional fixations at the start and end of the run. Subjects were instructed to attend to the film at all times, and to try to understand the original temporal and causal structures of the scrambled film. The stimuli were projected from a Propixx projector (VPixx Technologies, Bruno, Canada), with a resolution of 1920 pixels × 1080 pixels and a refresh rate of 60 Hz. The films were projected onto the center of the screen, with a 22.6° × 15.1° field of view. The background music that accompanied the film was delivered by MRI compatible in-ear headphones (MR Confon; Cambridge Research Systems, Rochester, UK).

### Image preprocessing

Structural images were bias field corrected and spatially normalized to the Montreal Neurological Institute (MNI) space. The first 10 images of the functional data were discarded to allow the MR signal to achieve T1 equilibration. Functional images were motion-corrected using the six rigid-body transformation parameters. After motion correction, there was no difference in the framewise displacement (FD)^56,57^ between the moments of low and high understanding, in both the Initial (high: FD = 0.039 ± 0.007, low: FD = 0.039 ± 0.007; *t*(66) = 0.511, *p* = 0.611) and Repeated (high: FD = 0.041 ± 0.006, low: FD = 0.041 ± 0.006; *t*(66) = 1.282, *p* = 0.204) Scrambled conditions. The functional images were slice timing-corrected, intensity-normalized, and registered to MNI-aligned T1-weighted images. We applied the FMRIB’s ICA-based X-noiseifier (FIX) to automatically identify and remove noise components.^58–60^ The BOLD time series was linearly detrended and band pass filtered (0.009 Hz < *f* < 0.125 Hz) to remove low frequency confounds and high frequency physiological noise. The data were spatially smoothed with a Gaussian kernel of full width at half maximum of 5 mm. All analyses were conducted in the volumetric space and the cortical surface of the MNI standard template was reconstructed using Freesurfer^61^ for visualization purposes.

### Whole brain parcellation

Cortical regions were parcellated into 114 ROIs following Yeo et al.^23^ based on a seven-network cortical parcellation estimated from the resting-state functional data of 1,000 adults.^24^ Subcortical regions were parcellated into eight ROIs, corresponding to the bilateral amygdala, hippocampus, thalamus, and striatum, extracted from the Freesurfer segmentation of the FSL MNI152 template brain.^23^ The time series of the voxels within each ROI were averaged, resulting in a matrix of functional scan duration (1870 s) × region (122 ROIs). To replicate the results using different atlases, we used the BNA that parcellates the whole brain into 246 ROIs without spatial overlap.^30^ For a comparison between the two parcellation schemes, we calculated the topological overlap between each BNA ROI and the eight pre-defined functional networks of Yeo et al,^24^ which was regarded as the probability of a specific BNA ROI being identified as part of each of the eight functional networks. The network label with the highest probability was assigned to each BNA ROI.

### Dynamic functional connectivity and network analysis

Sliding window correlation was used to measure dynamic FCs of the pairwise regions.^29,62–65^ A window size of 36 s was used, which matches the average duration of a scrambled scene of the film. The chosen window size fell within the range of optimal window size suggested by previous research.^66–68^ A tapered window, convolved with a Gaussian kernel of σ = 3 s, was used to give higher weights to the center of the window.^29,69,70^ An L1 penalty was added to increase the sparsity of the resulting correlation matrices, using the Graphical Lasso.^71^ The regularization parameter was fixed to λ= 0.01 for the ROIs selected from the Yeo et al.^24^ atlas and to λ = 0.1 for the BNA ROIs.^71^ The regularized correlation matrices were then Fisher’s *z*-transformed. Using sparse, weighted, and undirected FC matrices, we conducted graph theoretical network analyses, using the Brain Connectivity Toolbox (https://sites.google.com/site/bctnet/).^25^ As a global network measure, we calculated modularity by iteratively maximizing the modular structures using the Louvain algorithm^72–75^ with a resolution parameter *γ* = 1. Both the positive and negative edges were included, but a reduced weight was given to the negative edges, due to a higher significance of positive weights in the community structure of the functional brain.^14,76^ Further, we quantified global efficiency, after thresholding the matrices by leaving only the positive edges.^25^ The global efficiency was measured as the average inverse shortest path length between all pairs of regions in the network.^77^ Next, as regional measures of across-modular and within-modular connections, we calculated the participation coefficient and within-module degree z-score, based on the time-resolved community structure derived from the Louvain modularity algorithm (see Supplementary Method for details).^14,32^

For each subject, the time-resolved network measures were averaged to produce a single summary value representing either a moment of high or low level of understanding. The results from all subjects across three film stimuli were combined, and the Wilcoxon signed-rank tests were performed between the summary network measures that corresponded to high and low understanding moments. Statistical values from the regional network analysis and all subsequent analyses were FDR corrected for multiple comparisons across different functional networks.^78,79^ Additionally, we compared the FC strengths of the pairwise regions of the DMN. The ROIs indicated as part of the DMN from Yeo et al.^24^ were grouped into five regions – the mPFC, MFG, MTG, angular gyrus, and PreCu/PCC – based on their anatomical separations.^7^ The BOLD time series was extracted from each subregion of the DMN, and the Fisher’s r-to-z transformed correlation matrices, without L1 regularization, were calculated for all pairs using tapered sliding windows. Likewise, the average FC strengths corresponding to moments of high and low understanding were computed for each individual and compared using Wilcoxon signed-rank tests.

### Hidden Markov model

We defined ROIs based on group-level ICA,^39^ using FSL-MELODIC (http://www.fmrib.ox.ac.uk/fsl/melodic/index.html). The fMRI data of all subjects in all three conditions of the three different films were concatenated. The ICs were automatically extracted, then the authors hand-labeled the signal ICs, which resulted in total of 30 ICs. If an IC i) spatially overlapped with white matter or cerebrospinal fluid (CSF), ii) was derived from motion artifacts, or iii) had a temporal frequency that lied outside of a signal range (*f* > 0.125 Hz), it was discarded as a ‘noise’ IC. The signal ICs were qualitatively validated from their spatial overlaps with the well-known ICs identified from resting-state functional networks defined by Smith et al.^80^

We iteratively searched for the optimal number of states (K), within a range of two to eight. Since there is no straightforward, data-driven method of selecting the optimal K due to the known problem that the free energy of the model monotonically decreases as K increases,^38^ K was determined based on the model consistency^35^ and generalizability^6^ computed using the BOLD time series of the Original condition. We first tested the model consistency across iterations, where the same HMM training and inference procedures were repeated five times using the same hyper-parameters. The inferred sequences of latent states between pairwise iterations were compared. In addition, we tested the model generalizability, examining whether the model’s inferred sequence of the held-out subject’s fMRI responses could explain the average fMRI response patterns of other subjects who watched the same film. The assumption was that neural dynamics are synchronized across subjects who watch the same films.^81^ After K was determined based on the criteria of model consistency and generalizability among five iterations, we chose the one that had the highest log probability of the inferred state sequence, given the observed BOLD time series.

The HMM was trained using the concatenated time series of all subjects’ Original conditions of three different film datasets (hmmlearn.hmm.GaussianHMM). To overcome the problem of local minima during the initialization of the HMM inference, we initialized the HMM parameters using the output of k-means clustering with the same K (sklearn.cluster.KMeans). Expectation-Maximization (EM)^82^ of the forward-backward algorithm was used to estimate the optimal model parameters - transition probability, and emission probability. The log-likelihood of the observation was iteratively estimated, conditioned on the model. The number of iterations with different centroid seeds was set to 500. We decided that the forward-backward algorithm approached an asymptote when the gain in log-likelihood reached 0.001 during the re-estimation process. No restraint was given to the transition probability matrix so that the transitions could occur to all possible states. We modeled the emission probabilities using a mixture Gaussian density function, where the mean vector and covariance matrix were produced from the 30^th^ mixture components for each state. The mean vector was characterized as the weights given to the activation of the 30 ICs, and the covariance matrix was characterized as the functional covariance between the pairwise ICs.^35,36^ We defined each inferred neural state as the weighted sum of the extracted ICs with the mean vectors. To label each neural state as a known functional network, we masked the whole brain with Yeo et al.’s^24^ eight functional networks and compared the levels of activation corresponding to each network. The latent state was defined as a functional network that showed the highest level of activation. In instances where the two functional networks had comparable activation profiles, the state was termed using both networks (e.g., SM + VIS). The covariance matrix of each state was defined by the pairwise temporal covariance of the 30 ICs during the emergence of a discrete latent state within the fitted sequence. We applied the Louvain modularity algorithm to all latent states’ covariance matrices with varying Ks, and the output was consistently three modules. The modules were largely grouped as i) DMN + FPN, ii) VIS, and iii) SM, which were identified from the module’s probabilistic spatial correspondence to the resting-state functional networks defined by Smith et al.^80^

The estimated transition and emission probabilities were applied to decode the most probable sequence of the concatenated time series of all subjects during the Initial and Repeated Scrambled conditions, using the Viterbi algorithm.^83^ The outcome of the Viterbi algorithm is the probability of each latent state being most dominant at a specific time point. We chose the state with the highest probability to be a latent state at a specific moment, thus discretizing the latent sequence. To observe whether a neural state was dominantly associated with a certain cognitive state, we measured fractional occupancy during the moments of high and low understanding per subject. We conducted paired t-tests to compare fractional occupancy per state. We compared the pairwise-subject similarity of the inferred state sequences between the Initial and Repeated Scrambled conditions. For all pairwise subjects, the proportion of the paired time points having the identical latent state was measured. The state similarity measures were compared between the Initial and Repeated Scrambled conditions using a paired t-test.

### Dynamic predictive modeling

For the FC-pattern based decoding, the Fisher’s r-to-z transformed correlation matrices, without L1 regularization, were calculated for pairwise regions of 122 ROIs using tapered sliding windows. The FC strengths of each time window were used as features, corresponding to binary indices of understanding levels. Feature selection was employed in the FC pattern-based decoding. For the training dataset in each cross-validation fold, every pairwise region’s dynamic FC strength was correlated with changing levels of story understanding (Pearson’s correlation). If the distribution of training subjects’ correlations with the behavioral measure was significantly different from zero (one-sample t-test, *p* < 0.01), regardless of whether the average correlation was positive or negative, the feature was used in the SVM. In the activation-pattern based decoding, the time series of the 122 ROIs were used as features. For both types of decoding, we conducted normalization (*Z-*transformation) within each feature dimension per individual, thus maintaining within-feature temporal variance while removing across-subject and across-feature variances. A linear SVM (sklearn.svm.LinearSVC) was trained to classify the level of understanding (high vs. low), given the neural patterns at every time point of the training subjects. The model was tested on the held-out subject’s every time point and was cross-validated in a leave-one-subject-out fashion. The decoding accuracy was calculated by the proportion of correct classifications (high vs. low) over the total number of time points, which generates a chance level of 0.5. The performance in every cross-validation was averaged to represent an overall decoding performance. The actual decoding accuracy was compared to the distribution of decoding accuracies obtained from the permuted null data (*p* = (1 + number of null accuracies ≥ actual accuracy) / (1 + number of permutations), with number of permutations = 1,000). For permutation, the labels of high and low were randomly shuffled in every iterative training and testing session, while maintaining the distribution of consecutive labels of the nearby time points comparable to the actual behavioral index.

## Notes

### Competing Interest Statement

The authors have declared no competing interest.

## References

1. Langston, M. & Trabasso, T. (1999). Modeling causal integration and availability of information during comprehension of narrative texts. In van Oostendorp, H., & Goldman, S. R. (ed.) The Construction of Mental Representations during Reading. Lawrence Erlbaum Associates, Publishers. New Jersey: London.

2. Zwaan, R. A., Langston, M. C., & Graesser, A. C. (1995). The construction of situation models in narrative comprehension: An event-indexing model. Psychological Science 6, 292–297.

3. Graesser, A. C., Singer, M., & Trabasso, T. (1994). Constructing inferences during narrative text comprehension. Psychol. Rev. 101, 371–395.

4. Chang, C. H. C., Lazaridi, C., Yeshurun, Y., Norman, K. A., & Hasson, U. (2020). Relating the past with the present: Information integration and segregation during ongoing narrative processing. bioRxiv, https://doi.org/10.1101/2020.01.16.908731.

5. Chen, J., Leong, Y. C., Norman, K., & Hasson, U. (2017). Shared memories reveal shared structure in neural activity across individuals. Nat. Neurosci. 20(1): 115–25.

6. Baldassano, C., Chen, J., Zadbood, A., Pillow, J. W., Hasson, U., & Norman, K. A. (2017). Discovering Event Structure in Continuous Narrative Perception and Memory. Neuron 95, 709–721.

7. Simony, E., Honey, C. J., Chen, J., Lositsky, O., Yeshurun, Y., Wiesel, A., & Hasson, U. (2016). Dynamic reconfiguration of the default mode network during narrative comprehension. Nat. Commun. 7:12141.

8. Aly, M., Chen, J., Turk-Browne, N. B. & Hasson, U. (2018). Learning Naturalistic Temporal Structure in the Posterior Medial Network. J. Cogn. Neurosci. 30, 1345–1365.

9. Hasson, U., Yang, E., Vallines, I., Heeger, D. J. & Rubin, N. (2008). A Hierarchy of Temporal Receptive Windows in Human Cortex. J. Neurosci. 28, 2539–2550.

10. Lerner, Y., Honey, C. J., Silbert, L. J. & Hasson, U. (2011). Topographic Mapping of a Hierarchy of Temporal Receptive Windows Using a Narrated Story. J. Neurosci. 31, 2906–2915.

11. Honey, C. J., Thesen, T., Donner, T. H., Silbert, L. J., Carlson, C. E., Devinsky, O., Doyle, W. K., Rubin, N., Heeger, D. J., & Hasson, U. (2012). Slow Cortical Dynamics and the Accumulation of Information over Long Timescales. Neuron 76, 423–434.

12. Mar, R. A. (2004). The neuropsychology of narrative: story comprehension, story production and their interrelation. Neuropsychologia 42, 1414–1434.

13. Tononi, G., Sporns, O. & Edelman, G. M. (1994). A measure for brain complexity: relating functional segregation and integration in the nervous system. Proc. Natl. Acad. Sci. 91, 5033–5037.

14. Shine, J. M., Bissett, P. G., Bell, P. T., Koyejo, O., Balsters, J. H., Gorgolewski, K. J., Moodie, C. A., & Poldrack, R. A. (2016). The Dynamics of Functional Brain Networks: Integrated Network States during Cognitive Task Performance. Neuron, 92(2): 544–54.

15. Shine, J. M. Breakspear, M., Bell, P. T., Martens, K. A.` E., Shine, R., Koyejo, O., Sporns, O., & Poldrack, R. A. (2019). Human cognition involves the dynamic integration of neural activity and neuromodulatory systems. Nat. Neurosci. 22, 289–296.

16. Bullmore, E. & Sporns, O. (2012). The economy of brain network organization. Nat. Rev. Neurosci. 13, 336–349.

17. Zalesky, A., Fornito, A., Cocchi, L., Gollo, L. L. & Breakspear, M. (2014). Time-resolved resting-state brain networks. Proc. Natl. Acad. Sci. 111, 10341–10346.

18. Rabiner, L. R. & Juang, B. H. (1986). An introduction to hidden Markov models. IEEE ASSP Magazine 3, 4–16.

19. Danke, A. H., Wiley, J. (2017). What about False Insights? Deconstructing the Aha! Experience along Its Multiple Dimensions for Correct and Incorrect Solutions Separately. Front. Psychol. 20. doi: 10.3389/fpsyg.2016.02077.

20. Wolfe, M. B. W., Magliano, J. & Larsen, B. (2005). Causal and Semantic Relatedness in Discourse Understanding and Representation. Discourse Processes 39(2-3): 165–187.

21. Landauer, T. K. & Dumais, S. T. (1997). A solution to Plato’s problem: The latent semantic analysis theory of acquisition, induction, and representation of knowledge. Psychological Review 104, 211–240.

22. Sadaghiani, S., Poline, J.-B., Kleinschmidt, A. & D’Esposito, M. (2015). Ongoing dynamics in large-scale functional connectivity predict perception. Proc. Natl. Acad. Sci. 112, 8463–8468.

23. Yeo, B. T. T., Tandi, J. & Chee, M. W. L. (2015). Functional connectivity during rested wakefulness predicts vulnerability to sleep deprivation. NeuroImage 111, 147–158.

24. Yeo, B. T. T., Krienen, F. M., Sepulcre, J., Sabuncu, M. R., Lashkari, D., Hollinshead, M., Roffman, J. L., Smoller, J. W., Zöllei, L., Polimeni, J. R., Fischl, B., Liu, H., & Buckner, R. L. (2011). The organization of the human cerebral cortex estimated by intrinsic functional connectivity. J. Neurophysiol. 106(3): 1125–1165.

25. Rubinov, M. & Sporns, O. (2010). Complex network measures of brain connectivity: uses and interpretations. Neuroimage 52, 1059–1069.

26. Bassett, D. S., Porter, M. A., Wymbs, N. F., Grafton, S. T., Carlson, J. M., & Mucha, P. J. (2013). Robust detection of dynamic community structure in networks. Chaos 23, 013142.

27. Bullmore, E. & Sporns, O. (2009). Complex brain networks: graph theoretical analysis of structural and functional systems. Nat. Rev. Neurosci. 10, 186–198.

28. Achard, S. & Bullmore, E. (2007). Efficiency and cost of economical brain functional networks. PLoS Comput. Biol. 3, e17.

29. Allen, E. A. Damaraju, E., Plis, S. M., Eichele, T., & Calhoun, V. D. (2014). Tracking whole-brain connectivity dynamics in the resting state. Cereb. Cortex 24, 663–676.

30. Fan, L., Li, H., Zhuo, J., Zhang, Y., Wang, J., Chen, L., Yang, Z., Chu, C., Xie, S., Laird, A. R., Fox, P. T., Eickhoff, S. B., Yu, C., & Jiang, T. (2016). The Human Brainnetome Atlas: A New Brain Atlas Based on Connectional Architecture. Cereb. Cortex 26(8): 3508–26.

31. Xia, M., Wang, J. & He, Y. (2013). BrainNet Viewer: a network visualization tool for human brain connectomics. PLoS ONE 8, e68910.

32. Guimerà, R. & Nunes Amaral, L. A. (2005). Functional cartography of complex metabolic networks. Nature 433, 895–900.

33. Christoff, K., Irving, Z. C., Fox, K. C. R., Spreng, R. N. & Andrews-Hanna, J. R. (2016). Mind-wandering as spontaneous thought: a dynamic framework. Nat. Rev. Neurosci. 17, 718–731.

34. Andrews-Hanna, J. R., Reidler, J. S., Sepulcre, J., Poulin, R. & Buckner, R. L. (2010). Functional-Anatomic Fractionation of the Brain’s Default Network. Neuron 65, 550–562.

35. Vidaurre, D., Abeysuriya, R., Becker, R., Quinn, A. J., Alfaro-Almagro, F., Smith, S. M., & Woolrich, M. W. (2018). Discovering dynamic brain networks from big data in rest and task. NeuroImage 180, 1–11.

36. Vidaurre, D., Smith, S. M., & Woolrich, M. (2018). Brain network dynamics are hierarchically organized in time. Proc. Natl. Acad. Sci. 114(48): 12827–32.

37. Quinn, A. J., Vidaurre, D., Abeysuriya, R., Becker, R., Nobre, A. C., & Woolrich, M. W. (2018). Task-Evoked Dynamic Network Analysis Through Hidden Markov Modeling. Front. Neurosci., 12: 603.

38. Baker, A. P., Brookes, M. J., Rezek, I. A., Smith, S. M., Behrens, T., Smith, P. J. S., & Woolrich, M. (2014). Fast transient networks in spontaneous human brain activity. eLife 3: e01867.

39. Beckmann, C. F., DeLuca, M., Devlin, J. T., Smith, S. M. (2005). Investigations into resting-state connectivity using independent component analysis. Phil. Trans. R. Soc. B. 360, 1001–1013.

40. Shen, X., Finn, E. S., Scheinost, D., Rosenberg, M. D., Chun, M. M., Papademetris, X., & Constable, R. T. (2017). Using connectome-based predictive modeling to predict individual behavior from brain connectivity. Nat. Protoc. 12, 506–518.

41. Gonzalez-Castillo, J., Hoy, C. W., Handwerker, D. A., Robinson, M. E., Buchana, L. C., Saad, Z. S., & Bandettini, P. A. (2015). Tracking ongoing cognition in individuals using brief, whole-brain functional connectivity patterns. Proc. Natl. Acad. Sci. 112, 8762–8767.

42. Esterman, M., Rosenberg, M. D. & Noonan, S. K. (2014). Intrinsic fluctuations in sustained attention and distractor processing. J. Neurosci. 34, 1724–1730.

43. Rosenberg, M. D., Finn, E. S., Scheinost, D., Papademetris, X., Shen, X., Constable, R. T., & Chun, M. M. (2016). A neuromarker of sustained attention from whole-brain functional connectivity. Nat. Neurosci. 19, 165–171.

44. Honey, C. J., Newman, E. L. & Schapiro, A. C. (2018). Switching between internal and external modes: A multiscale learning principle. Netw. Neurosci. 1, 339–356.

45. Dixon, M. L., Fox, K. C. R. & Christoff, K. (2014). A framework for understanding the relationship between externally and internally directed cognition. Neuropsychologia 62, 321–330.

46. Chien, H.-Y. S., & Honey, C. J. (2020). Constructing and Forgetting Temporal Context in the Human Cerebral Cortex. Neuron 106, 1–12.

47. Franklin, N. T., Norman, K. A., Ranganath, C., Zacks, J. M., & Gershman, S. J. (2020). Structured Event Memory: A Neuro-Symbolic Model of Event Recognition. Psych. Rev., 127(3), 327–361.

48. Wehbe, L., Murphy, B., Talukdar, P., Fyshe, A., Ramdas, A., & Mitchell, T. (2014). Simultaneously Uncovering the Patterns of Brain Regions Involved in Different Story Reading Subprocesses. PLoS ONE 9, e112575.

49. Huth, A. G., de Heer, W. A., Griffiths, T. L., Theunissen, F. E. & Gallant, J. L. (2016). Natural speech reveals the semantic maps that tile human cerebral cortex. Nature 532, 453–458.

50. Vodrahalli, K., Chen, P. H., Liang, Y., Baldassano, C., Chen, J., Yong, E., Honey, C., Hasson, U., Ramadge, P., Norman, K. A., & Arora, S. (2018). Mapping between fMRI responses to movies and their natural language annotations. NeuroImage 180, 223–31.

51. Brainard, D. H. (1997). The Psychophysics Toolbox. Spatial Vision 10, 433–436.

52. Pelli, D. G. (1997). The VideoToolbox software for visual psychophysics: transforming numbers into movies. Spatial Vision 10, 437–442.

53. Zacks, J. M. & Swallow, K. M. (2007). Event Segmentation. Curr. Dir. Psychol. Sci. 16, 80–84.

54. Nishida, S. & Nishimoto, S. (2018). Decoding naturalistic experiences from human brain activity via distributed representations of words. NeuroImage 180, 232–242

55. Walther, D. & Koch, C. (2006). Modeling attention to salient proto-objects. Neural Networks 19, 1395–1407

56. Power, J. D., Barnes, K. A., Snyder, A. Z., Schlaggar, B. L. & Petersen, S. E. (2012). Spurious but systematic correlations in functional connectivity MRI networks arise from subject motion. NeuroImage 59, 2142–2154.

57. Power, J. D., Mitra, A., Laumann, T. O., Snyder, A. Z., Schlaggar, B. L., & Petersen, S. E. (2014). Methods to detect, characterize, and remove motion artifact in resting state fMRI. NeuroImage 84, 320–341.

58. Salimi-Khorshidi, G., Douaud, G., Beckmann, C. F., Glasser, M. F., Griffanti, L., & Smith, S. M. (2014). Automatic denoising of functional MRI data: combining independent component analysis and hierarchical fusion of classifiers. NeuroImage 90, 449–468.

59. Griffanti, L., Salimi-Khorshidi, G., Beckmann, C. F., Auerbach, E. J., Douaud, G., Sexton, C. E., Zsoldos, E., Ebmeier, K. P., Filippini, N., Mackay, C. E., Moeller, S., Xu, J., Yacoub, E., Baselli, G., Ugurbil, K., Miller, K. L., & Smith, S. M. (2014). ICA-based artefact removal and accelerated fMRI acquisition for improved resting state network imaging. NeuroImag, 95, 232–47.

60. Griffanti, L., Douaud, G., Bijsterbosch, J., Evangelisti, S., Alfaro-Almagro, F., Glasser, M. F., Duff, E. P., Fitzgibbon, S., Westphal, R., Carone, D., Beckmann, C. F., & Smith, S. M. (2017). Hand classification of fMRI ICA noise components. NeuroImage, 154: 188–205.

61. Fischl, B. (2012). Freesurfer. NeuroImage 62, 774–81.

62. Hutchison, R. M., Womelsdorf, T., Allen, E. A., Bandettini, P. A., Calhoun, V. D., Corbetta, M., Penna, S. D., Duyn, J. H., Glover, G. H., Gonzalez-Castillo, J., Handwerker, D. A., Keilholz, S., Kiviniemi, V., Leopold, D. A., de Pasquale, F., Sporns, O., Walter, M., & Chang, C. (2013). Dynamic functional connectivity: Promise, issues, and interpretations. NeuroImage 80: 10.1016.

63. Sakoğlu, Ü., Pearlson, G. D., Kiehl, K. A., Wang, Y. M., Michael, A. M., & Calhoun, V. D. (2010). A method for evaluating dynamic functional network connectivity and task-modulation: application to schizophrenia. MAGMA, 23(5-6): 351–66.

64. Handwerker, D. A., Roopchansingh, V., Gonzalez-Castillo, J., & Bandettini, P. A. (2012). Periodic changes in fMRI connectivity. NeuroImage, 63(3): 1712–9.

65. Leonardi, N., & Van De Ville, D. (2015). On spurious and real fluctuations of dynamic functional connectivity during rest. NeuroImage, 104: 430–6.

66. Liégeois, R., Ziegler, E., Phillips, C., Geurts, P., Gómez, F., Bahri, M. A., Yeo, B. T. T., Soddu, A., Vanhaudenhuyse, A., Laureys, S., & Sepulchre, R. (2016). Cerebral functional connectivity periodically (de)synchronizes with anatomical constraints. Brain Struct. Funct., 221(6): 2985–97.

67. Deng, L., Sun, J., Cheng, L., & Tong, S. (2016). Characterizing dynamic local functional connectivity in the human brain. Sci. Rep., 6: 26976.

68. Shirer, W. R., Ryali, S., Rykhlevskaia, E., Menon, V., & Greicius M. D. (2012). Decoding subject-driven cognitive states with whole-brain connectivity patterns. Cereb. Cortex, 22(1): 158–65.

69. Preti, M. G., Bolton, T. A. W., & Van De Ville, D. (2017). The dynamic functional connectome: State-of-the-art and perspectives. NeuroImage, 160: 41–54.

70. Barttfeld, P., Uhrig, L., Sitt, J. D., Sigman, M., Jarraya, B., & Dehaene, S. (2015). Signature of consciousness in the dynamics of resting-state brain activity. Proc. Natl. Acad. Sci. 112(3): 887–92.

71. Friedman, J., Hastie, T. & Tibshirani, R. (2008). Sparse inverse covariance estimation with the graphical lasso. Biostatistics 9, 432–441.

72. Newman, M. E. J. (2004). Fast algorithm for detecting community structure in networks. Phys. Rev. E. 69, 066133.

73. Newman, M. E. J. (2006). Modularity and community structure in networks. Proc. Natl. Acad. Sci. 103, 8577–8582.

74. Blondel, V. D., Guillaume, J.-L., Lambiotte, R. & Lefebvre, E. (2008). Fast unfolding of communities in large networks. J. Stat. Mech. 2008, P10008.

75. Fortunato, S. (2010). Community detection in graphs. Physics Reports 486, 75–174.

76. Rubinov, M. & Sporns, O. (2011). Weight-conserving characterization of complex functional brain networks. NeuroImage 56, 2068–2079.

77. Latora, V. & Marchiori, M. (2001). Efficient behavior of small-world networks. Phys. Rev. Lett. 87, 198701.

78. Benjamini, Y. & Hochberg, Y. (1995). Controlling the False Discovery Rate: A Practical and Powerful Approach to Multiple Testing. Journal of the Royal Statistical Society. Series B (Methodological) 57, 289–300.

79. Benjamini, Y. & Yekutieli, D. (2001). The control of the false discovery rate in multiple testing under dependency. Ann. Statist. 29, 1165–1188.

80. Smith, S. M., Fox, P. T., Miller, K. L., Glahn, D. C., Fox, M., Mackay, C. E., Filippini, N., Watkins, K. E., Toro, R., Laird, A. R., & Beckman, C. F. (2009). Correspondence of the brain’s functional architecture during activation and rest. Proc. Natl. Acad. Sci., 106(31): 13040–5.

81. Hasson, U., Nir, Y., Levy, I., Fuhrmann, G. & Malach, R. (2004) Intersubject Synchronization of Cortical Activity During Natural Vision. Science 303, 1634–1640.

82. Dempster, A. P., Laird, N. M. & Rubin, D. B. (1977). Maximum Likelihood from Incomplete Data via the EM Algorithm. Journal of the Royal Statistical Society. Series B (Methodological) 39, 1–38.

83. Rezek, I. & Roberts, S. (2005). Ensemble Hidden Markov Models with Extended Observation Densities for Biosignal Analysis. in Probabilistic Modeling in Bioinformatics and Medical Informatics (eds. Husmeier, D., Dybowski, R. & Roberts, S.), Springer, 419–450.

