## Supplementary Information for "Cognitive and Neural State Dynamics of Story Comprehension"

(A) Initial Scrambled condition

High > Low Understanding

| | Size | x | y | z | Region |
|---|---|---|---|---|---|
| 1 | 793 | -3 | +66 | +33 | Right Precuneus, Posterior Cingulate |
| 2 | 227 | -51 | +72 | +30 | Right Angular Gyrus |
| 3 | 202 | +39 | +84 | +39 | Left Precuneus, Posterior Cingulate |
| 4 | 134 | -3 | -51 | +39 | Right Medial Frontal Gyrus |
| 5 | 98 | -27 | -36 | +42 | Right Middle Frontal Gyrus |
| 6 | 56 | +9 | -63 | -15 | Left Medial Frontal Gyrus |
| 7 | 40 | +69 | +21 | -12 | Left Middle Temporal Gyrus |

Low > High Understanding

| | Size | x | y | z | Region |
|---|---|---|---|---|---|
| 1 | 1257 | +24 | +93 | +9 | Left Occipital |
| 2 | 560 | -12 | +96 | +3 | Right Occipital |
| 3 | 119 | +39 | +6 | +51 | Left Precentral Gyrus |
| 4 | 48 | -54 | +57 | +3 | Right Middle Temporal Gyrus |

(B) Repeated Scrambled condition

High > Low Understanding

| | Size | x | y | z | Region |
|---|---|---|---|---|---|
| | | | - | None | - |

Low > High Understanding

| | Size | x | y | z | Region |
|---|---|---|---|---|---|
| 1 | 183 | +24 | +96 | +9 | Left Occipital |
| 2 | 123 | -21 | +93 | +9 | Right Occipital |
| 3 | 48 | +42 | +81 | -18 | Left Fusiform Gyrus |

**Supplementary Table S1**. Results of general linear model analysis in the Initial and Repeated Scrambled conditions. The size (number of voxels) and the coordinates of the peak voxel of each activated cluster in the Montreal Neurological Institute space are reported, which correspond to Figure 3A (A) and Figure 3B (B).

| Latent states | Functional Networks | | | | | | | |
|---|---|---|---|---|---|---|---|---|
| | FPN | DMN | DAN | VAN | VIS | SM | LIMB | SUBC |
| SM+VIS | -3.42 | -4.00 | -0.21 | 2.33 | 3.57 | 5.76 | -0.59 | 1.31 |
| DAN | 1.16 | -3.46 | 10.15 | 1.88 | -8.67 | 1.41 | -0.30 | -0.00 |
| Integrated DMN+VIS | 0.75 | 2.14 | -1.09 | -1.55 | 1.79 | -3.07 | 0.18 | -1.00 |
| Segregated DMN+VIS | 1.07 | 4.27 | -8.34 | -1.94 | 2.69 | -2.63 | 0.61 | 0.14 |

**Supplementary Table S2**. Average blood oxygen-level dependent (BOLD) activations in eight functional network masks in relation to the four inferred latent neural states. The activation maps of the latent states were defined by the weighted sum of the 30 independent components with the mean vector of the inferred emission probabilities (mixture Gaussian). The eight pre-defined functional network masks were applied to the four latent neural states and the average BOLD responses of the voxels within each mask were calculated. The label was assigned as the functional network that showed a maximum degree of activation. If more than one functional network showed a comparably high degree of activation, then both networks were used to label a corresponding latent state. DAN: dorsal attention network, DMN: default mode network, FPN: frontoparietal network, LIMB: limbic network, SM: somatosensory-motor network, SUBC: subcortical networks, VAN: ventral attention network, VIS: visual network.

| | FC | | | | Activation |
| --- | --- | --- | --- | --- | --- |
| | Within-film decoding | | Across-film decoding | | |
| | Initial | Repeated | Initial | Repeated | |
| Whole brain | 528.2 ± 104.0 | 146.8 ± 17.4 | 776.5 ± 142.7 | 142.1 ± 13.1 | 122 |
| FPN | 190.4 ± 56.2 | 48.9 ± 8.9 | 303.2 ± 60.0 | 49.0 ± 12.4 | 26 |
| DMN | 190.0 ± 22.0 | 68.0 ± 30.2 | 280.8 ± 36.0 | 61.0 ± 8.6 | 26 |
| DAN | 192.8 ± 66.8 | 35.0 ± 6.9 | 264.9 ± 87.2 | 34.1 ± 16.9 | 14 |
| VAN | 158.0 ± 23.7 | 42.2 ± 6.6 | 246.0 ± 40.2 | 41.6 ± 7.1 | 24 |
| VIS | 100.1 ± 35.5 | 30 ± 8.4 | 130.8 ± 27.5 | 35.1 ± 12.4 | 10 |
| SM | 64.7 ± 27.1 | 16.3 ± 6.4 | 105.6 ± 21.1 | 19.4 ± 7.6 | 10 |
| LIMB | 22.7 ± 13.4 | 5.3 ± 2.2 | 27.5 ± 11.8 | 4.0 ± 2.5 | 4 |
| SUBC | 42.3 ± 10.3 | 16.9 ± 6.1 | 56.5 ± 8.5 | 48.4 ± 9.9 | 8 |

**Supplementary Table S3**. The number of features used in predictive models. The selected number of functional connectivities was lesser in the Repeated Scrambled condition than in the Initial Scrambled condition. The additional feature selection was not applied to activation pattern-based decoding owing to the initial small number of regions of interest used as features. DAN: dorsal attention network, DMN: default mode network, FPN: frontoparietal network, LIMB: limbic, SM: somatosensory-motor, SUBC: subcortical networks, VAN: ventral attention, VIS: visual network.

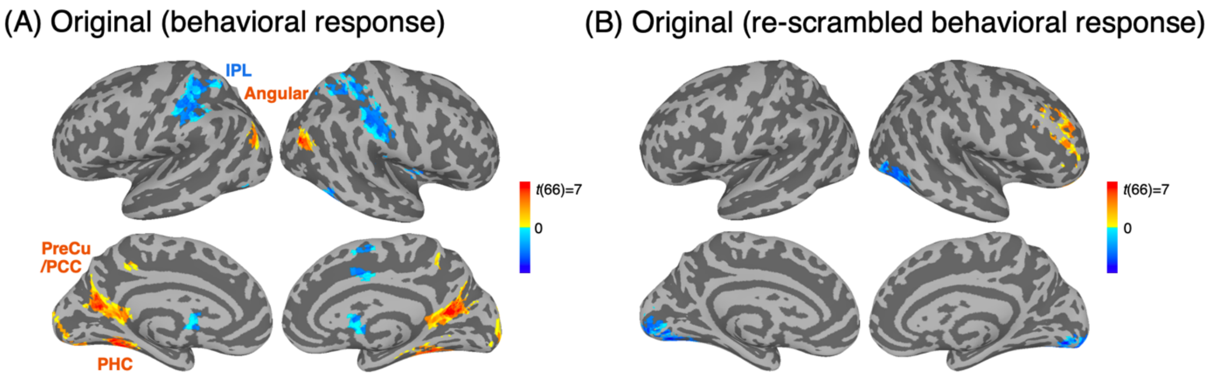


**Supplementary Figure S1**. Modulation of blood oxygen-level dependent (BOLD) responses in the Original condition (cluster size > 40 and $q < 0.01$). (A) Contrast between the moments of high and low understanding, defined by behavioral responses while subjects were watching films in an intact sequence (Original condition). The results were comparable to those of the Initial Scrambled condition (Figure 3A). During the moments of high understanding, the default mode network, including the bilateral precuneus (PreCu) and posterior cingulate (PCC) [-15 +57 +15], and bilateral angular gyrus (Angular) [-45 +78 +27] [+39 +75 +30] exhibited higher BOLD responses. The bilateral inferior parietal lobe (IPL) [-54 +21 +39] [+42 +33 +42], a part of the dorsal attention network, and the visual sensory network [+48 +72 -12] increased their BOLD responses when the understanding was low. Interestingly, during the moments of high understanding in the Original condition, increases in BOLD responses were also found in the bilateral parahippocampal cortex (PHC) [-30 +48 -9] [+30 +51 -9]. The PHC, which was previously considered a part of the contextual association network along with the PreCu and PCC, is known to increase its BOLD response when the given objects are congruent with the scene contexts, compared to when they are incongruent.[1-3] In the Original condition, it is possible that subjects were able to incorporate their additional understanding within the constructed narrative contexts, which they previously had failed to comprehend during the Initial Scrambled condition. (B) Contrast between the moments physically matching the moments of high and low understanding of the Scrambled conditions. We re-scrambled the sequence of the behavioral index extracted during viewing of the scrambled films in order of the original film sequence. There was no notable modulation of the BOLD responses, indicating that the results of Figure 3A were not likely driven by the intrinsic properties of the film stimuli.

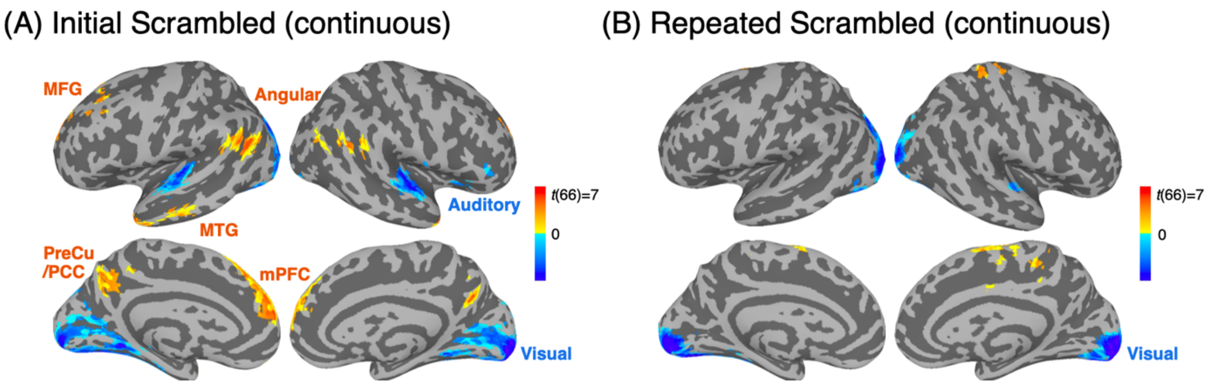


**Supplementary Figure S2**. Modulation of blood oxygen-level dependent (BOLD) responses by changes in the level of story understanding when a continuous behavioral index was used as a regressor (cluster size > 40 and $q < 0.01$). (A) Regions that showed positive and negative correlations with a behavioral index of understanding in the Initial Scrambled condition. The results are consistent with Figure 3, showing larger BOLD responses in the default mode network as the understanding increases, which includes the left middle frontal gyrus (MFG) [+8 -34 +52] and medial prefrontal cortex (mPFC) [+15 -60 +24], the bilateral middle temporal gyrus (MTG) [-45 -18 -42] [+57 -3 -33], the bilateral angular gyrus (Angular) [+54 +66 +27] [-63 +51 +36], and the precuneus/posterior cingulate (PreCu/PCC) [0 +63 +36]. When the understanding is low, the sensory processing regions, including the visual (Visual) [+30 +78 -15] and auditory (Auditory) [+48 +21 +6] [-51 +12 +3] cortices, exhibited higher BOLD responses. (B) Regions that showed positive and negative correlations with the behavioral index in the Repeated Scrambled condition. The results are similar to Figure 3, with no significant modulation of the BOLD responses, except in early visual areas [-18 +93 +0].

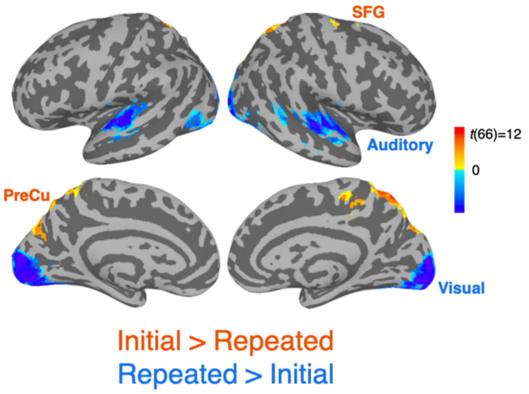


**Supplementary Figure S3**. Modulation of blood oxygen-level dependent (BOLD) responses across the two Scrambled conditions (cluster size > 40 and $q < 0.01$). During the Initial Scrambled condition, BOLD responses in the bilateral precuneus (PreCu) [-6 +69 +66] and right superior frontal gyrus (SFG) [-27 +6 +72] increased compared to the Repeated Scrambled condition. The two regions, included in the frontoparietal network, are known to be involved in goal-directed thoughts and cognitive control.[4,5] During the Repeated Scrambled condition, the visual (Visual) [+3 +87 -3] and auditory (Auditory) [-69 +18 +3] [+48 +18 +6] cortices showed higher BOLD responses compared to the Initial Scrambled condition. The low-level sensory processing regions were more dominant when the same film was watched repeatedly, which may be due to comparatively increased sensory processing when subjects are cognitively disengaged after having already understood the story.

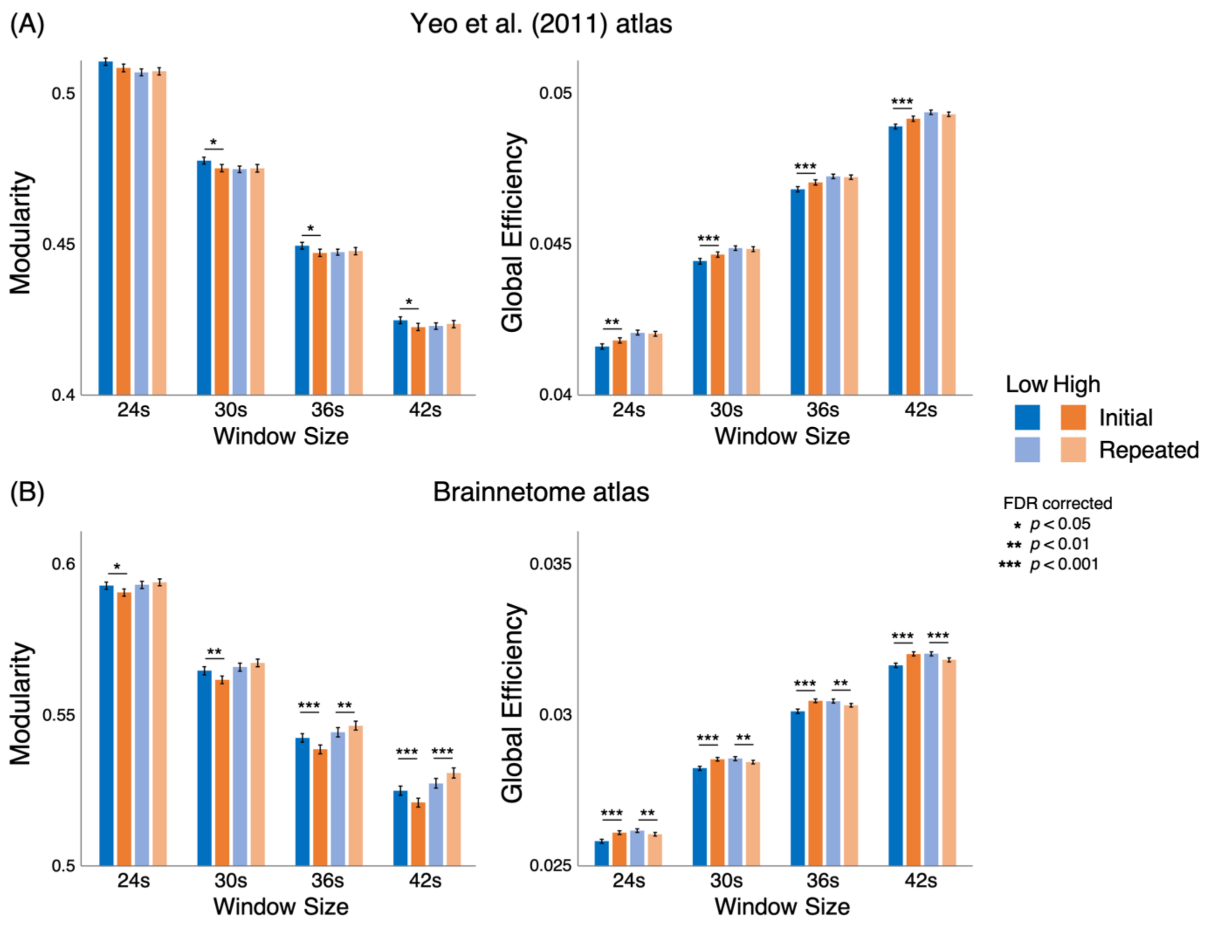


**Supplementary Figure S4**. Global network measures across four different window sizes (24, 30, 36, and 42s). The atlas of (A) Yeo et al. (2011) (122 ROIs) and (B) the Brainnetome atlas (246 ROIs) were used. The results are comparable to Figure 4B. (A) Modularity significantly decreased during the moments of high story understanding, only in the Initial Scrambled condition (all FDR-*ps* < 0.05 except window size 24s, FDR-*p* = 0.065), but not in the Repeated Scrambled condition (all FDR-*ps* > 0.5). Likewise, global efficiency increased during the moments of high understanding, only in the Initial Scrambled condition (all FDR-*ps* < 0.01), but not in the Repeated Scrambled condition (all FDR-*ps* > 0.4). We observed a significant interaction between the Scrambled conditions (Initial and Repeated) and story understanding levels (high and low) for all window sizes, for both modularity (all *ps* < 0.05) and efficiency (all *ps* < 0.01). There was a main effect of varying window sizes (all *ps* < 0.001). (B) As in (A), modularity significantly decreased during the moments of high understanding in the Initial Scrambled condition (all FDR-*ps* < 0.05), but not in the Repeated Scrambled condition (FDR-*ps* < 0.15 for window sizes of 24 s and 30 s, but FDR-*ps* < 0.01 for window sizes of 36 s and 42 s). Likewise, global efficiency increased during the moments of high understanding in the Initial Scrambled condition (all FDR-*ps* < 0.001). In the Repeated Scrambled condition, the efficiency showed significant decrease during the moments of high understanding (all FDR-*ps* < 0.01). We observed a main effect of window size (all $p$s < 0.001) as well as a significant interaction between the Scrambled conditions and story understanding levels for all window sizes, for both modularity (all $p$s < 0.01) and efficiency (all $p$s < 0.001).

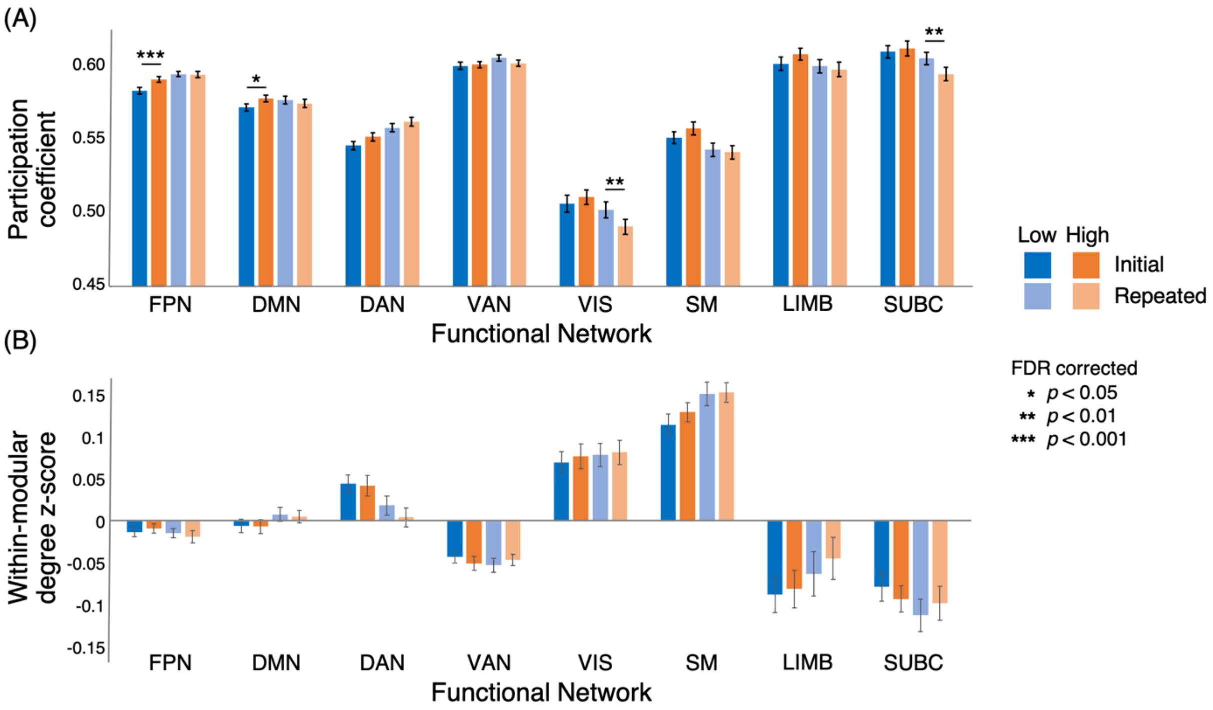


**Supplementary Figure S5**. Regional network measures (window size 36s), using 122 regions of interest. (A) Participation coefficient. The across-modular functional connectivities (FCs) in the frontoparietal network and the default mode network increased when the understanding was high, only in the Initial Scrambled but not in the Repeated Scrambled condition, with a significant interaction between the Scrambled conditions and understanding levels. (B) Within-modular degree z-score. Within-modular FCs did not vary in any of the functional networks depending on the level of understanding. DAN: dorsal attention network, DMN: default mode network, FPN: frontoparietal network, LIMB: limbic network, SM: somatosensory-motor networks, SUBC: subcortical networks, VAN: ventral attention network, VIS: visual network.

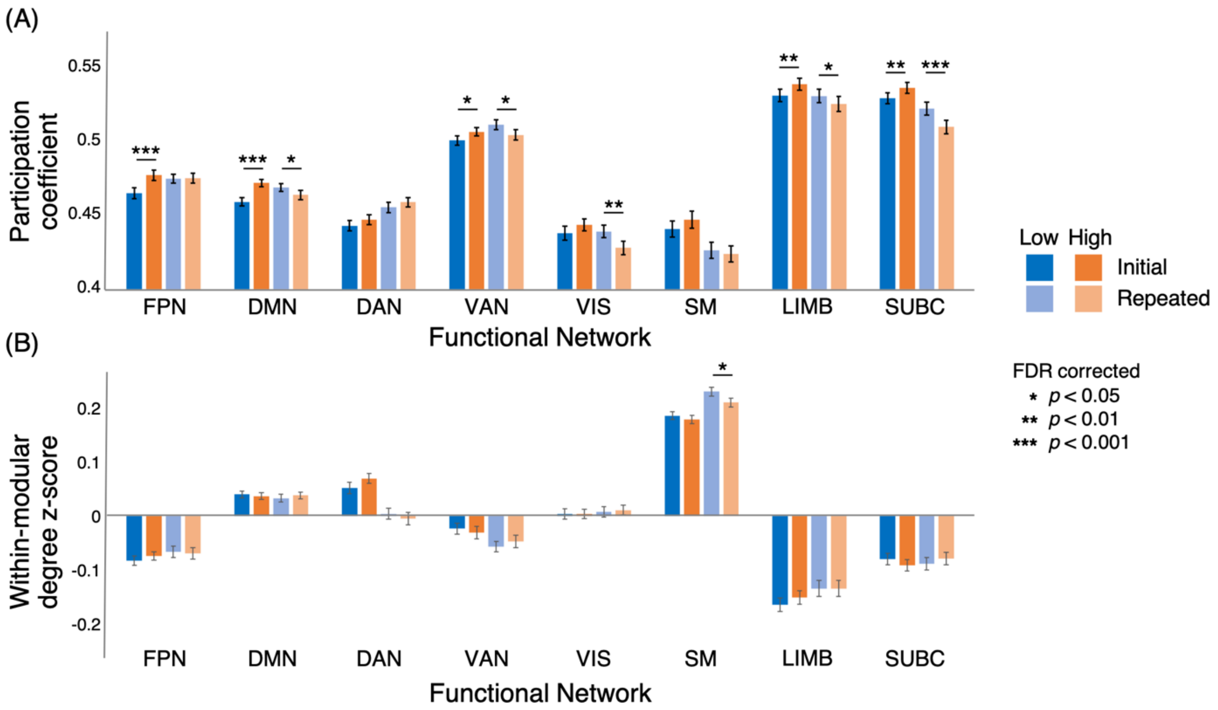


**Supplementary Figure S6**. Regional network measures (window size of 36 s), using 246 regions of interest (ROIs) of the Brainnetome atlas. (A) Participation coefficient. All functional networks exhibited trends of increase in participant coefficients during the moments of high understanding, only in the Initial Scrambled condition. The frontoparietal network ($z(66) = 4.110$) and default mode network ($z(66) = 5.422$) showed significantly higher participation coefficients (all FDR-$ps < 0.001$), with a significant interaction effect (FPN: $F(1,66) = 10.79$, DMN: $F(1,66) = 42.99$; all $ps < 0.01$). (B) Within-modular degree z-score. None of the functional networks modulated the WMDZ as the level of story understanding changed across time. DAN: dorsal attention network, DMN: default mode network, FPN: frontoparietal network, LIMB: limbic network, SM: somatosensory-motor networks, SUBC: subcortical networks, VAN: ventral attention network, VIS: visual network.

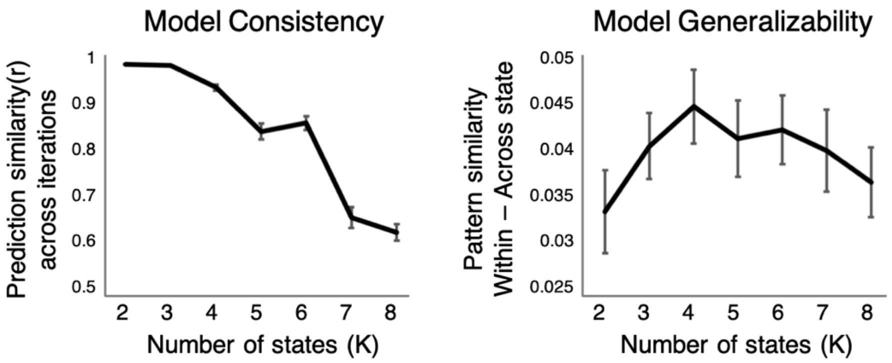


**Supplementary Figure S7**. The hyperparameter selection (K; number of discrete states), based on the model consistency and generalizability. Model consistency indicates the stability of the model's predicted latent state sequence across repeated iterations using the same hyper-parameters. The inference procedures were repeated five times, and the average of the pairwise sequence similarities was calculated to assess the consistency of the model. Model generalizability measures if the model trained on an individual subject's functional magnetic resonance imaging (fMRI) response could explain the average fMRI response patterns of the rest of the subjects who watched the same film. K of 4 and 6 were chosen to be the optimal number of states, given their high consistencies and generalizability among possible Ks.

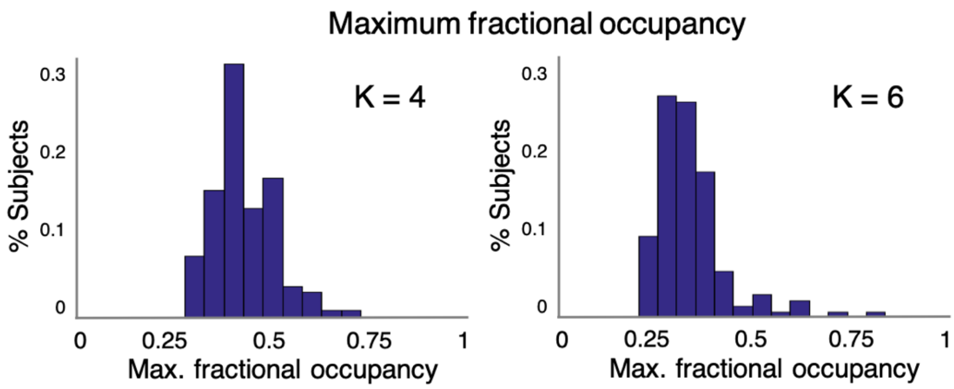


**Supplementary Figure S8**. Validation of the dynamics in the inferred sequence from the hidden Markov model (HMM). The fractional occupancy of the highest emerging state was calculated per subject's inferred sequence. The maximum fractional occupancy was less than 50% for most subjects (*p*s < 0.001 for both Ks), indicating that the HMM captured the dynamics of state transitions.

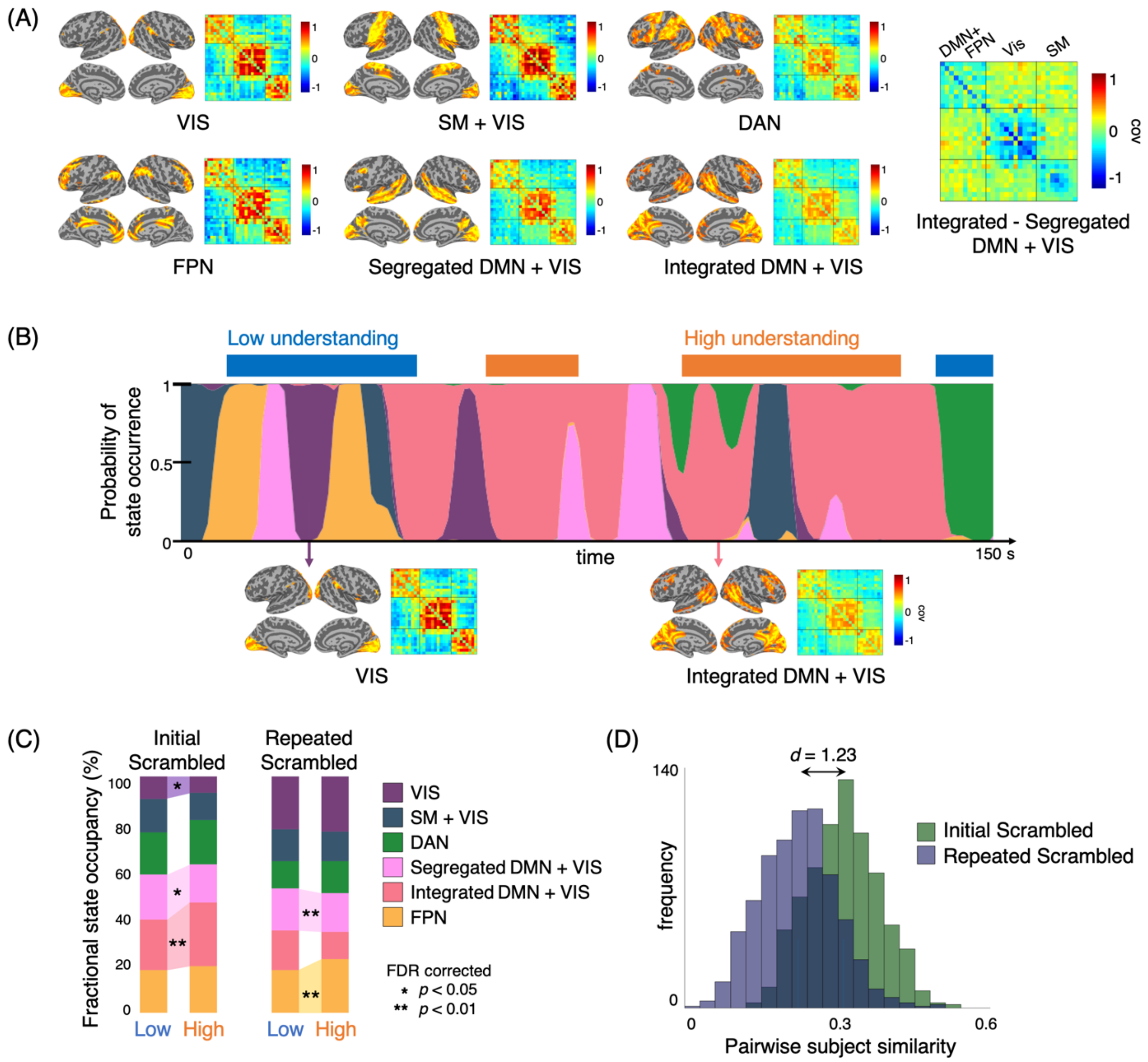


**Supplementary Figure S9**. Results of the hidden Markov model inference using K = 6. (A) The mean activation and functional covariance of the six latent states. We observed the visual sensory network and frontoparietal network in addition to the four states shown in Figure 5A. (B) The state occupancy and transition dynamics of a representative subject in relation to the cognitive states of understanding, during the Initial Scrambled condition. (C) The average fractional occupancies of the six latent states during moments of high and low understanding. In the Initial Scrambled condition, an Integrated DMN + VIS had a higher fractional occupancies during high, compared to low, understanding ($t(66) = 3.57$, FDR-$p < .01$). The VIS had a lower fractional occupancy during low understanding ($t(66) = 2.85$, FDR-$p < .05$). In the Repeated Scrambled condition, we observed a significant decrease in the fractional occupancy of Segregated DMN + VIS ($t(66) = 3.86$, FDR-$p < .01$), as well as an increase in the fractional occupancy of FPN ($t(66) = 33.3$, FDR-$p < .01$), during moments of high, compared to low, understanding. (D) Synchrony of the latent neural state across individuals, in the Initial and Repeated Scrambled conditions. The across-subject similarity was higher in the Initial, compared to the Repeated Scrambled condition ($t(718) = 25.90$, $p < 0.001$, Cohen's $d = 1.23$). DAN: dorsal attention network, DMN: default mode network, FPN: frontoparietal network, SM: somatosensory-motor networks, VIS: visual network.

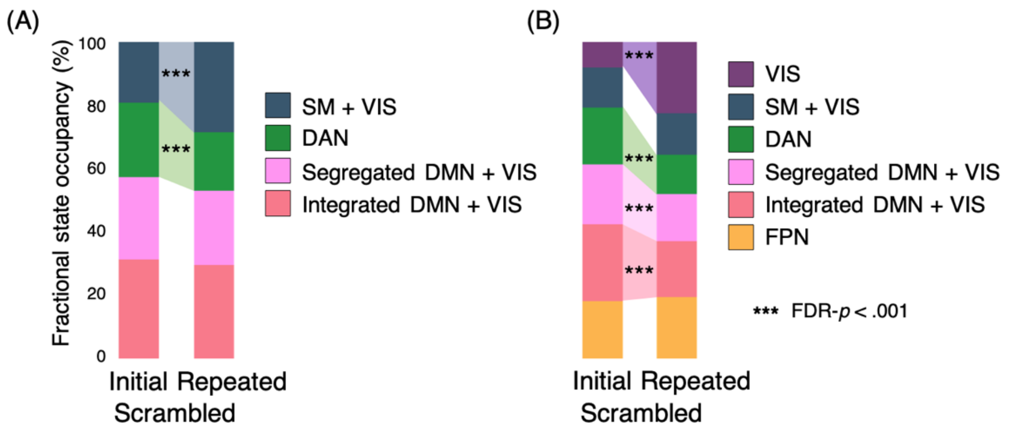


**Supplementary Figure S10**. Comparison of the fractional occupancy of states between the Initial and Repeated Scrambled conditions, in (A) K = 4 and (B) K = 6. (A) When K of 4 was used, there was a higher fractional occupancy of the DAN in the Initial Scrambled condition ($t(66) = 6.48$, FDR-$p < .001$), whereas the SM + VIS had a higher fractional occupancy during the Repeated Scrambled condition ($t(66) = 5.42$, FDR-$p < .001$). Both Integrated and Segregated DMN + VIS network states showed a trend of a higher occupancy during the Initial compared to Repeated Scrambled condition, though not to a significant degree (FDR-$p < .125$). (B) When K of 6 was used, the two DMN + VIS had higher fractional occupancy during the Initial Scrambled condition ($t(66) = 6.37$ for Integrated, $t(66) = 4.38$ for Segregated DMN + VIS, both FDR-$p$s $< 0.001$). Similarly, the DAN had a higher level of occupancy during the Initial Scrambled condition ($t(66) = 9.13$, FDR-$p < .001$). On the contrary, the VIS had a higher occupancy during the Repeated Scrambled condition ($t(66) = 6.95$, FDR-$p < .001$). Overall, the results are in line with our general linear model results that compared the Initial and Repeated Scrambled conditions (Supplementary Figure S3). Functional networks involved with higher-level cognition, such as the default mode network and the dorsal attention network, were more active when subjects were actively engaged in understanding a novel story. Lower-level sensory networks, such as the visual sensory network or the somatosensory-motor network, were highly involved when subjects were repeatedly watching the same scrambled films. DAN: dorsal attention network, DMN: default mode network, FPN: frontoparietal network, SM: somatosensory-motor networks, VIS: visual sensory network.

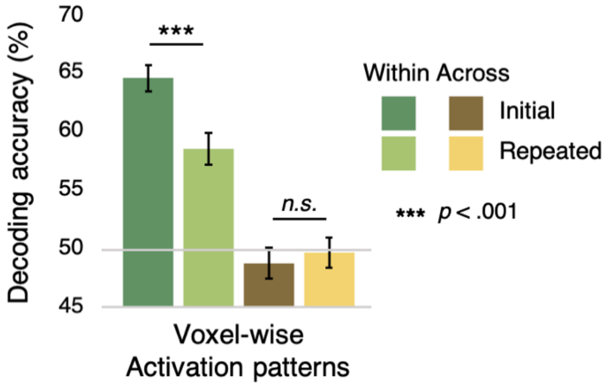


**Supplementary Figure S11**. Voxel-wise activation pattern-based prediction of the understanding level. In the within-film decoding, the performance of the Initial Scrambled condition was significantly higher than the Repeated Scrambled condition ($z(66) = 3.40$, $p < 0.001$). However, in the across-film decoding, there was no significant difference between the two conditions ($z(66) = 0.693$, $p = 0.4881$), comparable to the result of region of interest activation pattern-based decoding in Figure 7D. Two-way analysis of variance showed a significant interaction effect between the Scrambled condition (high or low) and the type of cross-validation method (within- or across-film) ($F(1,66) = 8.787$, $p < 0.01$). The number of features used in the within-film decoding was 5296 ± 659 for the Initial Scrambled condition and 2991 ± 214 for the Repeated Scrambled condition. In the across-film decoding, the selected number of features was 6046 ± 1797 for the Initial Scrambled condition and 4295 ± 157 for the Repeated Scrambled condition. The selected numbers of features were all higher than the numbers of selected functional connectivities, indicated in Supplementary Table S3.

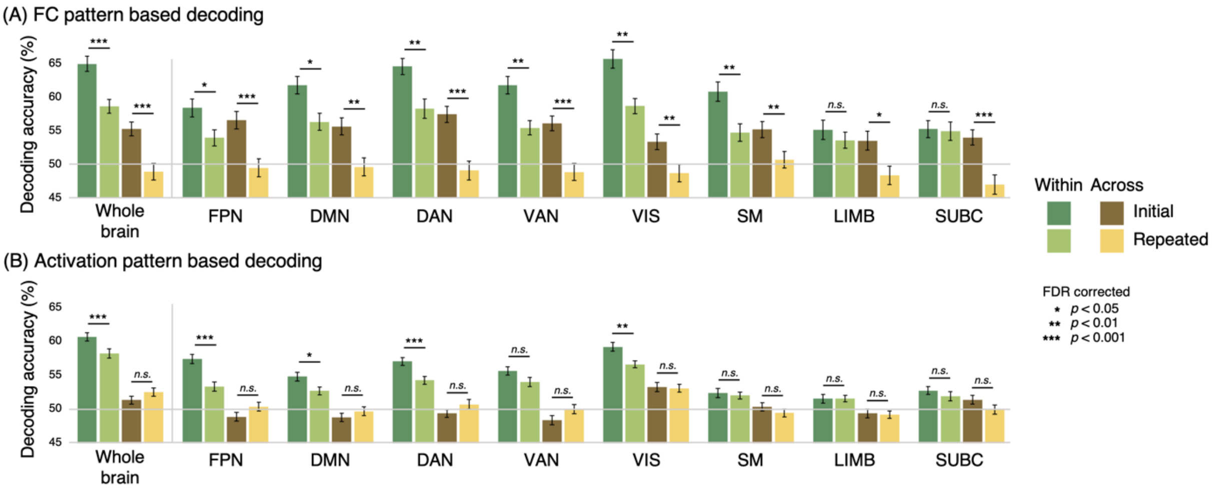


**Supplementary Figure S12.** Prediction of the understanding level using individual functional networks as neural features. (A) Functional connectivity (FC) pattern-based decoding. Among the FCs significantly correlated with the evolving level of story understanding, we selected FCs that have a particular functional network as seed regions. All individual functional networks showed significantly higher across-film prediction accuracy in the Initial compared to Repeated Scrambled condition (all FDR-*p*s < 0.05), which is comparable to the decoding results using whole brain FCs (Figure 7D). (B) Activation pattern-based decoding. In the across-film decoding, no significant difference between the Initial and Repeated Scrambled conditions was found in any of the functional networks (all FDR-*p*s > 0.127), which is comparable to the decoding results using whole brain ROIs (Figure 7D).

**References**


1. Bar, M. & Aminoff, E. (2003). Cortical Analysis of Visual Context. *Neuron* **38**, 347–358.
2. Kveraga, K., Ghuman, A. S., Kassam, K. S., Aminoff, E. A., Hämäläinen, M. S., Chaumon, M., & Bar, M. (2011). Early onset of neural synchronization in the contextual associations network. *Proc. Natl. Acad. Sci.* **108**, 3389–3394.
3. Lavenex, P. & Amaral, D. G. (2000). Hippocampal-neocortical interaction: a hierarchy of associativity. *Hippocampus* **10**, 420–430.
4. Niendam, T. A., Laird, A. R., Ray, K. L., Dean, Y. M., Glahn, D. C., & Carter, C. S. (2012). Meta-analytic evidence for a superordinate cognitive control network subserving diverse executive functions. *Cogn Affect Behav Neurosci* **12**, 241–268.
5. Spreng, R. N., Stevens, W. D., Chamberlain, J. P., Gilmore, A. W. & Schacter, D. L. (2010). Default network activity, coupled with the frontoparietal control network, supports goal-directed cognition. *Neuroimage* **53**, 303–317.
