## Supplementary Methods for "Cognitive and Neural State Dynamics of Story Comprehension"

### Global network measures: modularity and global efficiency

The calculation of modularity follows Newman[1] and Rubinov & Sporns[2], using undirected, weighted FC matrices with reduced weights given to the negative edges.

$$Q_T = \frac{1}{v^+}\Sigma_{ij}(w_{ij}^+ - e_{ij}^+)\,\delta_{M_iM_j} - \frac{1}{v^+ + v^-}\Sigma_{ij}(w_{ij}^- - e_{ij}^-)\,\delta_{M_iM_j} \quad (1)$$

Equation (1) indicates a time-resolved Louvain modularity algorithm ($Q_T$). $w_{ij}^+$ indicates the weights of the positive FCs between regions $i$ and $j$ within the range (0,1), and $w_{ij}^-$ indicates the weights of the negative FCs between regions $i$ and $j$ within the range (0,1). $v^\pm$ indicates sum of all positive or negative connection weights within the graph, where $v^\pm$ equals $\Sigma_{ij}w_{ij}^\pm$. $\delta_{M_iM_j}$ indicates the module partitions between regions $i$ and $j$, where $\delta_{M_iM_j}=1$ identifies that $i$ and $j$ lie within the same module, and $\delta_{M_iM_j}= 0$ identifies that $i$ and $j$ lie in different modules. $e_{ij}^\pm$ indicates the strength of a connection divided by the total weight of the graph, where $e_{ij}^\pm = \frac{\Sigma_j w_{ij}^\pm\,\Sigma_i w_{ij}^\pm}{v^\pm}$.

$$E_{gT} = \frac{\Sigma_{i\neq j\in G}(d_{ij}^{\,w})^{-1}}{\mathrm{N(N-1)}} = \frac{1}{\mathrm{N(N-1)}}\sum_{i\neq j\in G}\frac{1}{d_{ij}^{\,w}} \quad (2)$$

Global efficiency is calculated based on Latora & Marchiori,[3] using a thresholded matrix of positive edges alone. The global efficiency was measured as the average inverse shortest path length between all pairs of regions in the network. Equation (2) is the measure of global efficiency ($E_{gT}$), where $d_{ij}^{\,w}$ indicates the shortest path length between the regions $i$ and $j$, and N indicates the total number of regions in the graph.

### Regional network measures: participation coefficient and within-module degree z-score

Based on the time-resolved community structure derived from the Louvain modularity algorithm, we calculated the time-resolved participation coefficients and the within module degree z-score measures for every ROI, following Guimerà & Nunes Amaral[4] and Shine et al.[5]

$$PC_{iT} = 1 - \sum_{s=1}^{N_{MT}}\left(\frac{\kappa_{isT}}{\kappa_{iT}}\right)^2 \quad (3)$$

$$WMDZ_{iT} = \frac{\kappa_{iT} - \acute{\kappa}_{SiT}}{\sigma_{\kappa_{SiT}}} \quad (4)$$

Equations (3) and (4) show the time-resolved measure of participation coefficient ($PC_{iT}$) and within-module degree z-score ($WMDZ_{iT}$). In (3), $PC_{iT}$ ranges between 0 to 1. $\kappa_{isT}$ indicates the strength of positive FCs between region $i$ and all other regions at module $S_i$ at time T, and $\kappa_{iT}$ indicates the strength of positive FCs of region $i$ to all other regions, irrespective of module assignment. $N_{MT}$ indicates the number of modules at time T, where the modules were defined using the Louvain algorithm. The time-resolved participation coefficient of a region approximates to 1 if the connections are made with the regions of other modules. In (4), $\kappa_{iT}$ indicates the strength of connections of region $i$ to other regions that lie within the same module $S_i$ at time T, and $\acute{\kappa}_{SiT}$ indicates the average of $\kappa$ over all regions in the module $S_i$ at time T. $\sigma_{\kappa_{SiT}}$ indicates the standard deviation of $\kappa$ in module $S_i$ at time T.

**References**


1. Newman, M. E. J. (2006). Modularity and community structure in networks. *Proceedings of the National Academy of Sciences* **103**, 8577–8582.
2. Rubinov, M. & Sporns, O. (2011). Weight-conserving characterization of complex functional brain networks. *NeuroImage* **56**, 2068–2079.
3. Latora, V. & Marchiori, M. (2001). Efficient behavior of small-world networks. *Phys. Rev. Lett.* **87**, 198701.
4. Guimerà, R. & Nunes Amaral, L. A. (2005). Functional cartography of complex metabolic networks. *Nature* **433**, 895–900.
5. Shine, J. M., Bissett, P. G., Bell, P. T., Koyejo, O., Balsters, J. H., Gorgolewski, K. J., Moodie, C. A., & Poldrack, R. A. (2016). The Dynamics of Functional Brain Networks: Integrated Network States during Cognitive Task Performance. *Neuron* **92**(2), 544-54.
